# Excitatory-Inhibitory Recurrent Dynamics Produce Robust Visual Grids and Stable Attractors

**DOI:** 10.1101/2022.03.28.486063

**Authors:** Xiaohan Zhang, Xiaoyang Long, Sheng-Jia Zhang, Zhe Sage Chen

## Abstract

Spatially modulated grid cells has been recently found in the rat secondary visual cortex (V2) during activation navigation. However, the computational mechanism and functional significance of V2 grid cells remain unknown, and a theory-driven conceptual model for experimentally observed visual grids is missing. To address the knowledge gap and make experimentally testable predictions, here we trained a biologically-inspired excitatory-inhibitory recurrent neural network (E/I-RNN) to perform a two-dimensional spatial navigation task with multisensory (e.g., velocity, acceleration, and visual) input. We found grid-like responses in both excitatory and inhibitory RNN units, and these grid responses were robust with respect to the choices of spatial cues, dimensionality of visual input, activation function, and network connectivity. Dimensionality reduction analysis of population responses revealed a low-dimensional torus-like manifold and attractor, showing the stability of grid patterns with respect to new visual input, new trajectory and relative speed. We found that functionally similar receptive fields with strong excitatory-to-excitatory connection appeared within fully connected as well as structurally connected networks, suggesting a link between functional grid clusters and structural network. Additionally, multistable torus-like attractors emerged with increasing sparsity in inter- and intra-subnetwork connectivity. Finally, irregular grid patterns were found in a convolutional neural network (CNN)-RNN architecture while performing a visual sequence recognition task. Together, our results suggest new computational mechanisms of V2 grid cells in both spatial and non-spatial tasks.

**Highlights:**

- Grid patterns emerge in trained RNNs with multisensory inputs
- Grid patterns are robust to the RNN input and network connectivity
- Population responses show emergent ring-like manifolds and attractors
- Grid-like patterns persist in RNNs while performing a non-spatial task.

## INTRODUCTION

The discoveries of periodic grid cells or grid-like responses have been reported in the rat, mouse, bat and human brains during various spatial and non-spatial tasks (Hafting et al. 2005; Fyhn et al. 2007; Yartsev et al. 2011; Jacobs et al. 2013; Doeller et al. 2010; Constantinescu et al. 2016; Bellmund et al. 2018a; Bellmund et al. 2018b; Nau et al. 2018; Bao et al. 2019; Shilnikov and Maurer 2016; Rueckemann et al. 2021; Ginosar et al. 2021; Grieves et al. 2021). One of important roles of grid cells is to integrate self-motion information that provides a path integrative input to identify the spatial location even when external sensory inputs are lacking or noisy (Bush et al. 2014). Grid patterns were first found in single neurons of the rat medial entorhinal cortex (mEC) (Hafting et al. 2005; Fyhn et al. 2007), and recently have been reported in the rat primary somatosensory cortex (S1) (Long and Zhang 2021) and the rat secondary visual cortex (V2) (Long et al. 2021a). These S1 and V2 grid cells share some common features as mEC grid cells, such as conjunctive grid-head direction tunings and theta-modulated firing; furthermore, these grid-like responses are not disrupted from the absence of vibrissae or visual input (Long and Zhang 2021; Long et al. 2021a).

Attractor dynamics have been suggested in the hippocampal and entorhinal representations of the local environment (Wills et al. 2005; Bruak 2014; Agmon and Burak 2020). To date, many computational models of mEC grid cells have been proposed (for reviews, see (Giocomo et al. 2011; Rowland et al. 2016)), including the continuous attractor models (Fuhs and Touretzky 2006; Bruak and Fiete 2009), oscillator interference models (Burgess et al. 2007; Burgess 2008), feedforwad neural network with excitatory and inhibitory synaptic plasticity (Weber and Sprekeler 2018), and other hybrid models (Bush and Burgess 2014; Kang and Balasubramanian 2019; Rosay et al. 2019). Recent work has shown that grid cells emerge from trained RNNs that predict spatial location based on a pure velocity input (Banino et al. 2018; Cueva and Wei 2018; Sorscher et al. 2020), which supports the hypothesis of recurrent attractor dynamics and path integrator in the cognitive map (McNaughton et al. 2006), such that the attractor state may encode a stable representation of a variable (such as position) in the absence of external input.

Vision plays an important role in spatial navigation, and various visual cues can be integrated with spatial cues to guide movement. In the literature, spatial tunings have been established in the dorsal lateral geniculate nucleus (dLGN), primary visual cortex (V1) and other visual cortical areas from head-fixed or freely-foraging animals during spatial navigation tasks (Fiser et al. 2016; Hok et al. 2018; Saleem et al. 2018; Campbell and Giocomo 2018; Fournier et al. 2020; Diamanti et al. 2021; Flossmann and Rochefort 2021; Zong et al. 2022). Vision and movement jointly contribute to hippocampal place codes (Chen et al. 2013); simultaneous recordings of rodent V1 and hippocampal CA1 have also shown coherent coding of spatial signals (Ji and Wilson 2007; Haggerty and Ji 2015; Fournier et al. 2020).

To help understand the role of vision in forming grid responses, we developed a biologically-constrained recurrent neural network (RNN) to model the experimentally observed grid cells in the rat V2. We adapted our computational models to incorporate both visual and spatial cues in the RNN in a spatial navigation task. We investigated the impact of various spatial (velocity or acceleration) and visual cues (illuminance or optical flow) on the grid responses of the excitatory and inhibitory units. At the population level, we employed dimensionality reduction to reveal low-dimensional ring attractor dynamics, and investigated the stability of grid responses with respect to visual and spatial cues, synaptic connectivity and E/I balance. In parallel to the spatial navigation task, we also trained a combined convolutional neural network (CNN)-RNN model to perform a pure visual task and investigated the emerged grid patterns. Together, these simulation results suggest new computational mechanisms of grid cells in the visual cortex upon receiving multisensory input, and produce experimentally testable hypotheses for future investigation.

## RESULTS

### Trained RNNs Produced Robust Grid Patterns with Varying Spatial and Visual Cues

We trained biologically-constrained excitatory-inhibitory (E/I) RNNs (Song et al. 2016; Rajakumar et al. 2021; Xue et al. 2022) to perform a spatial navigation task in a two-dimensional (2D) environmental enclosure. We envisioned that the RNN received various forms of visual and spatial cues in the input (Table 1), and predicted the position. The network consisted of both excitatory and inhibitory units according to a 4:1 ratio (STAR Methods), and employed a non-negative rectified linear unit (ReLu) in the activation function (Figure 1A). We adopted a similar computation simulation setup to train the RNN (Banino et al. 2018; Cueva and Wei 2018; Sorscher et al. 2020), with additionally imposed constraints (e.g., Dale’s principle and clustered synaptic connections).

**Table 1:**
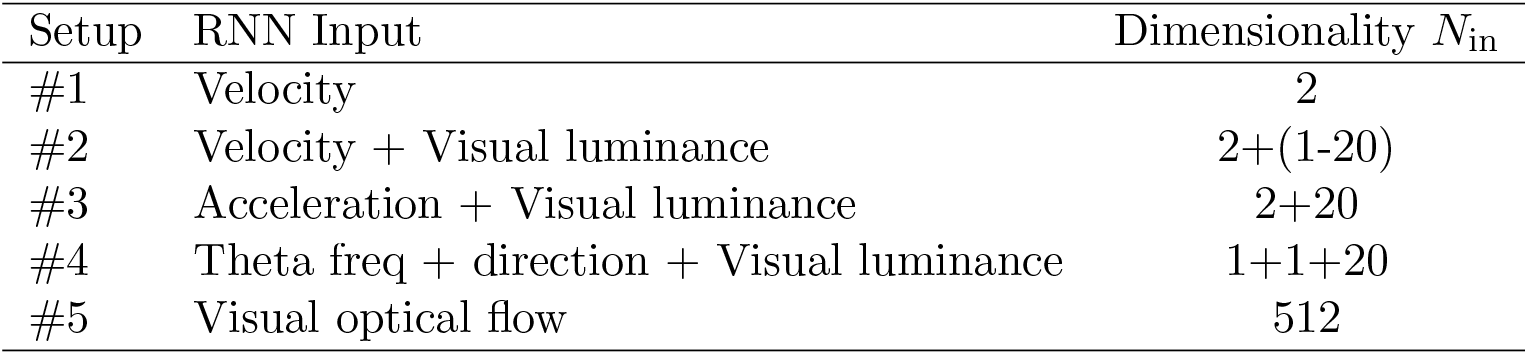
Testing various input configurations in the E/I-RNN.

| Setup | RNN Input | Dimensionality $N_{\text{in}}$ |
| --- | --- | --- |
| #1 | Velocity | 2 |
| #2 | Velocity + Visual luminance | 2+(1-20) |
| #3 | Acceleration + Visual luminance | 2+20 |
| #4 | Theta freq + direction + Visual luminance | 1+1+20 |
| #5 | Visual optical flow | 512 |

**Figure 1.**
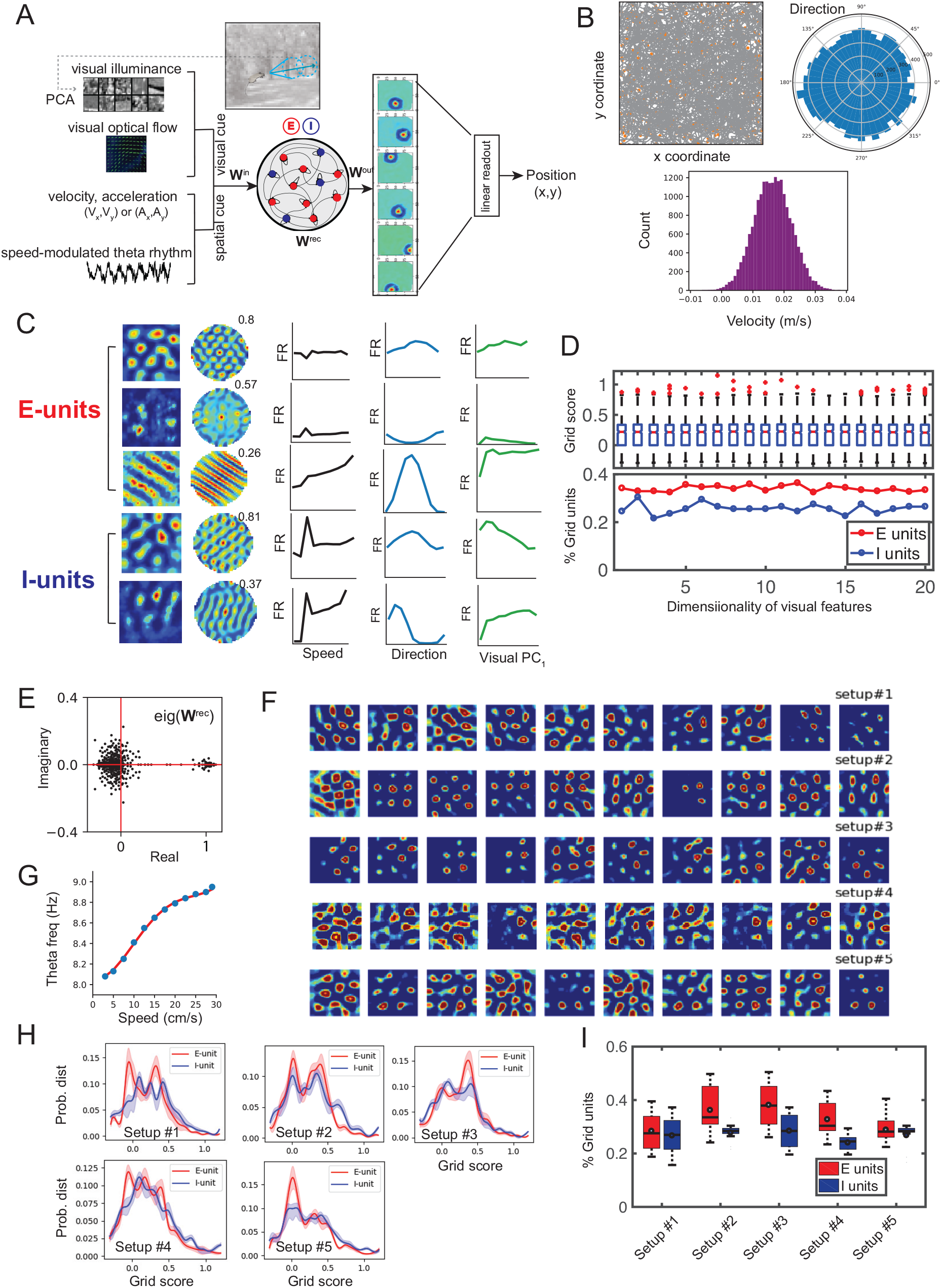
Multisensory Input and Recurrent Dynamics of E/I-RNN Produce Robust Grid Patterns. **(A)** Schematic of the excitatory-inhibitory (E/I)-RNN to perform a 2D spatial navigation task with multisensory cues. The visual cue may appear in the form of principal components (PCs) for the 8 *×* 8 image patch along the heading angle, or the form of visual optical flow. The spatial cue may appear in the form of velocity (speed and direction), acceleration, or speed-modulated theta rhythms. **(B)** Top left: Simulated trajectories (gray curve). Orange dots denote the place cell centers uniformly distributed within the 2D enclosure. Top right and bottom: Distributions of simulated run speed and direction statistics. **(C)** Examples of emerged grid-like and band-like patterns from excitatory and inhibitory units of the trained RNN (Setup #2). First column: firing rate (FR) heat map; second column: spatial autocorrelogram (the numbers indicate the grid score); third column: speed tuning curve; fourth column: direction tuning curve; fifth and sixth columns: tuning curve with respect to visual illumination principal component (PC_1_). All tuning curve panels are in the same scale (a.u.). **(D)** Statistics of grid units were relatively stable with respect to the dimensionality of visual features in PCA subspace (1-20). **(E)** Complex eigenspectrum of **W**^rec^ from a trained E/I-RNN. **(F)** Emergent grid-like RNN units with highest grid scores under different input configurations (Rows 1-5 corresponded to results from Setup #1-5). **(G)** The relationship between theta frequency and run speed, redrawn from Dannenberg et al., 2020). **(H)** Distributions of grid scores from the excitatory and inhibitory units under different input configurations. In each setup, statistics were generated from 10 trained RNNs. The shaded area along the curve represents *±* SD. **(I)** Percentages of grid units in the trained RNNs under different input configurations. Box plot statistics were generated from 10 trained RNNs in each setup. See also Figure S1.

#### RNN input configuration

First, to replicate previous computational simulation results (Banino et al. 2018; Cueva and Wei 2018; Sorscher et al. 2020), we employed the E/I-RNN with a pure velocity (i.e., speed and direction) input (Setup #1). The run speed followed a normal distribution and run direction was uniformly distributed between 0 and 360° (Figure 1B). Next, we added additional visual illuminance input with varying dimensionality (see STAR Methods, Setup #2). Upon convergence, we projected the hidden unit activations onto the 2D space to obtain the position-modulated, speed-modulated, direction-modulated and illumination-based tunings (Figures 1C and S1). We have witnessed a wide range of heterogeneity in spatially-tuning of the RNN units under different input configurations (Figure S1). Depending on specific configurations, subsets (20-50%) of excitatory and inhibitory units showed clear grid-like responses. The ranges of grid scores varied depending on the input configurations or cell types. These grid-like units displayed varying spatial frequencies as shown in their autocorrelograms, and also displayed conjunctive coding for the head direction and speed (Figure 1C). In addition to grid-like excitatory and inhibitory units, we also found some periodic band-like excitatory units (Figures 1C and S1), which appeared as a combination of multiple grid units (Krupic et al. 2012; Narvatilova et al. 2016). In Setup #2, we systematically varied the dimensionality of principal components of visual illumination features and found consistent grid patterns in the RNN units (Figure 1D). To examine the stability of the learned RNN, we calculated the eigenvalues of recurrent weight matrix **W**^rec^, and showed that a large majority of complex (or real) eigenvalues were within the unit circle (Figure 1E), whereas a very small percentage of eigenvalues were slightly greater than 1. This result may suggest the chaotic spontaneous activity present in the trained RNN (Sussillo and Abbott 2009). Furthermore, similar grid responses were observed when we replaced velocity (*V*_*x*_, *V*_*y*_) with acceleration 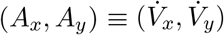 in the spatial input (Setup #3, Figure 1F). Motivated by the recent experimental data of theta-modulated firing in V2 grid cells (Long et al. 2021a), as well as the finding that the the theta frequency or amplitude increased proportion to the animal’s run speed (Chen et al. 2016; Dannenberg et al. 2020), we relaxed the assumption of direct speed access and used the frequency of theta oscillations as the input (Setup #4, Figure 1G). Consequently, we still observed robust emerged grid responses in the hidden units (Figure 1F). The results of Setup #3 and Setup#4 were not really surprising as these substituted variables involved only linear (or approximately linear) operations from velocity or speed. Notably, the direction input to the RNN was crucial to the formation of grid patterns, grid units did not emerge if we removed the direction in Setup #4.

Finally, we computed the optical flow cue from the consecutive visual scenes and use that vector fields as the input (see STAR Methods, Setup #5) and repeated the RNN training procedure. With the pure visual cue, the trained RNN still preserved the spatially-modulated grid patterns (Figure 1F). This result was also easy to interpret since the optical flow offered an indirect source of motion cue (i.e., direction and speed information). Overall, varying input configurations in our computer simulations yielded robust grid patterns with comparable grid score statistics (Figures 1H and 1I). In the remaining paper, we will focus the analyses on two configurations (Setup #2 and Setup #5).

#### Impact of the sequence length

The E/I-RNN was trained by batches of simulated trajectories with a fixed length; we found that a wide range of sequence lengths, *ℓ*, produced grid patterns from the trained RNN units (Figure 2A). Specifically, we varied *ℓ* from 5 to 30 (corresponding to 100-600 ms for the choice of 20-ms temporal bin size) and found that the GS statistics was robust with respect to the sequence length (Figure 2B). Our simulation results from multiple independently trained RNNs have shown the minimum *ℓ* that the achieved good GS statistic was 100 ms, roughly matching the timescale of one theta (5-10 Hz) cycle.

**Figure 2:**
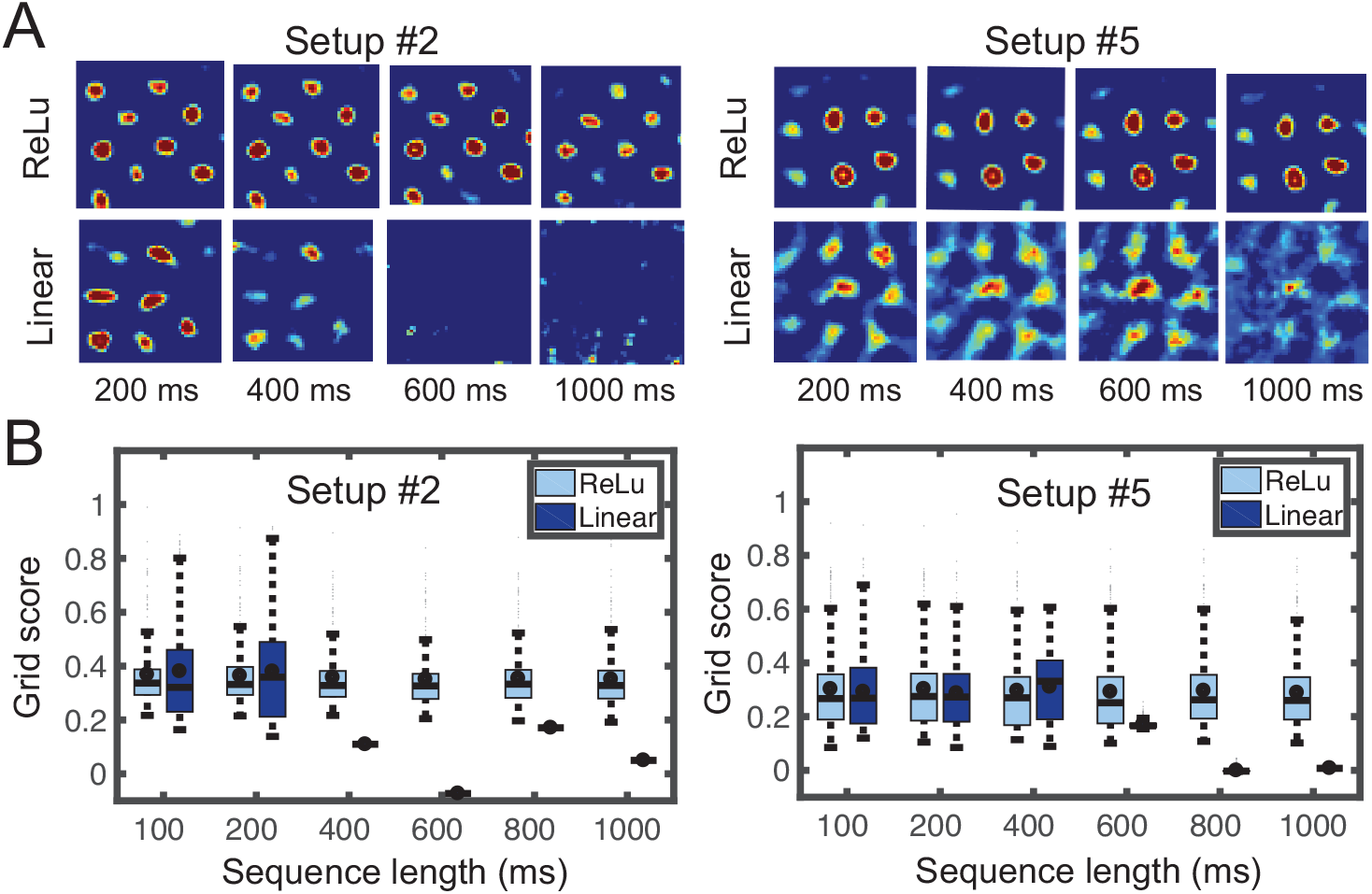
Impact of Sequence Length on Grid Patterns in Trained RNNs. **(A)** Grid-like patterns were robust with respect to a wide range of sequence lengths (200 ms, 400 ms, 600 ms, and 1000 ms) and activation functions in the trained E/I-RNN. **(B)** Grid score statistics with different sequence lengths. In each condition, top 50% of grid scores were used for better visualization. See also Figure S2.

#### Activation function

We further tested whether the relaxation of default ReLu (non-negativity) activation function to an unconstrained linear unit could yield similar results (Setup #2 and Setup #5). Interestingly, the trained linear E/I-RNN could still produce grid-like responses (Figure 2A), but the grid score was lower and the grid patterns were sensitive to the sequence length (Figure 2B). In this special case, the linear attractor network (without the non-negativity constraint) is a linear path integrator, integrating velocity (or acceleration) information in time to predict the future position (see Supplemental Information).

Symmetry breaking is a critical condition for complex pattern formation (McNaughton et al. 2006). It has been shown that the non-negativity of activation function suffices to generate the grid patterns (Sorscher et al. 2020); our results showed that even in the absence of non-negativity, Dale’s principle alone may be sufficient for symmetry breaking. As a sanity check, we also trained a RNN without the Dale’s principle constraint, fewer grid patterns emerged, but there was no clustered structure in the 2D embedding space (Figure S2C).

#### Mixed selectivity, paired unit correlation and emerged functional clusters

For each recurrent unit, we computed the grid score (GS), and the unit with GS*>*0.3 was empirically categorized as grid units. We then computed the percentage of grid cells from both excitatory and inhibitory populations. Among the identified grid units, we plotted their tuning curves with respect to direction, speed, acceleration, or theta frequency (Figures 1B and S1). In Setup #2, many of identified grid cells showed strong speed (71%) or directional (79%) tuning, or both (65%). Because of the high-dimensionality of visual features, we only plotted the tunings with respect to the dominant principal components (PCs) (e.g., Figure 1C). Speed tuning or directional tuning could also be observed for non-grid or band-like units (Figure S1). Interestingly, we found that a subset of band-like patterns had cosine-shaped direction tunings, all of which covered uniformly between 0° and 360° (Figure S3).

In Setup #2, we examined the impact of change in the speed and visual input on the grid field (Figure 3A). To examine the co-dependency of spatial and visual tunings for the *j*-th unit, we correlated the time-averaged unit firing rates contributed by spatial or visual input alone, 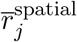 (by setting the visual input to zero) and 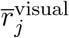 (by setting the velocity input to zero). We found statistically significant correlation between them (Figure 3B, left panel, Pearson’s correlation, *P* < 10^−5^), suggesting that main driving factor of mean firing rate was the internal recurrent dynamics instead of external visual or speed input in the standard setting. However, increasing the speed by 10 fold would substantially increased the mean firing rate (Figure 3B, right panel). Additionally, by setting the velocity to zero, we plotted the temporal firing rates of grid cells and measured the pairwise firing rate correlation when receiving a series of stimuli input (analogous to watching image sequences in a head-fixed setting). In some examples, we found that units with spatially similar grid fields showed temporally correlated visual responses (e.g., *E*_1_ vs. *E*_2_ units in Figure 3C); whereas in other examples, units with correlated grid fields showed uncorrelated visual responses (e.g., *E*_3_ vs. *E*_4_ units in Figure 3C), suggesting the independence between spatial and visual tunings. In a general setting, the excitatory grid units that had mutually strong excitatory synaptic connections tended to have similar grid fields; however, the converse was not necessarily true for the inhibitory grid units (for illustrated examples, see Figure 3D). Therefore, these functionally clustered grid units emerged as a result of strong synaptic connections, whereas weakly coupled grid units tended to be functionally decoupled.

**Figure 3:**
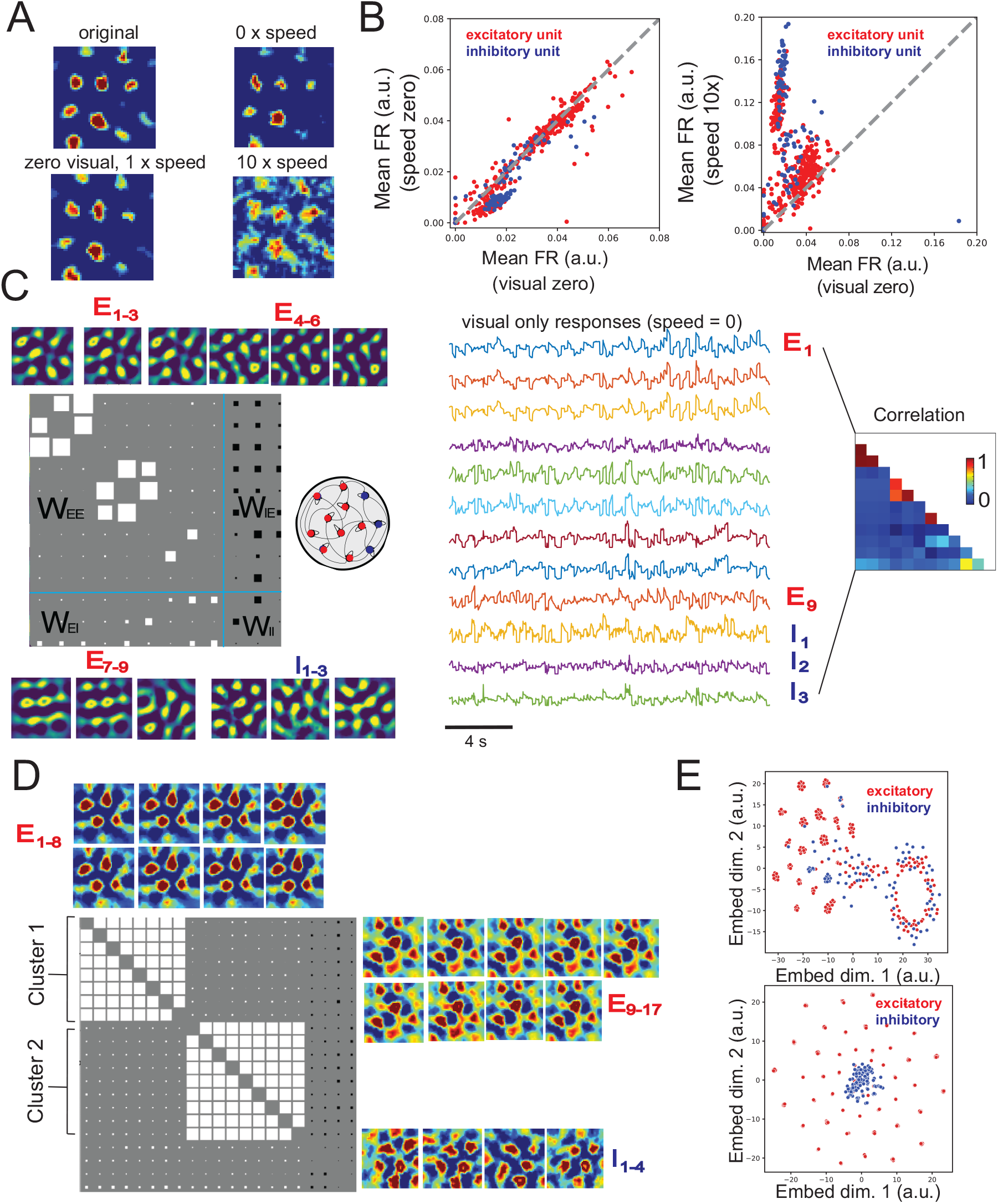
Mixed Selectivity of RNN units and Emerged Functional Clusters. **(A)** Examples of paired grid-like units and their co-activated firing with respect to visual input. **(B)** Scatter plots of 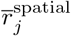 and 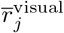 among the E/I-RNN units (Setup #2). Red and blue color denote excitatory and inhibitory units, respectively. **(C)** From a trained E/I RNN (Setup #2), selected 12 (9 excitatory plus 3 inhibitory) grid units and their weight connectivity (the white/black square denotes positive/negative synaptic weight, the size of square is proportion to the strength). Temporal traces represent the firing rates of these 12 units when setting velocity to zero. The lower triangular matrix denotes the correlation between 12 firing rate traces. **(D)** From a trained linear E/I-RNN (Setup #2), selected 21 (17 excitatory plus 4 inhibitory) grid units and their weight connectivity. **(E)** 2D embedding of RNN grid fields that were associated with top 60% GS (Setup #2, top: ReLu activation function; bottom: linear activation function). Red and blue color denote excitatory and inhibitory grid units, respectively. See also Figures S3 and S4.

To identify the functionally similar clusters in the grid cell subnetwork, we applied the stochastic neighborhood embedding (SNE) techniques to embed the grid fields onto a 2D space for visualization. Interestingly, we found grid-like patterns form many distinct clusters in the embedded space, especially among the excitatory grid-like units; this observation was robust regardless the chosen activation function or setup (Figures 3E and S4).

### Low-dimensional Ring Manifolds and Attractors Emerged from Trained E/I-RNNs

Next, we examined the neural representation of high-dimensional population activity of E/I-RNN units. According to the percentage of the explained variance (Figure 4A), we visualized two of the first three dominant PCs onto the latent space. We found an emergent 2D ring-shaped manifold in the PC subspace (Figure 4B). In Setup #2, when the simulated sequence length was short, the PC_1_-PC_2_ plane formed a ring attractor primarily explained by the dominant spatial components; in contrast, when the sequence length was very long, the ring attractor was occupied on the PC_2_-PC_3_ plane. This was possibly because the visual input of the longer visuospatial sequence contributed to more variance of RNN’s hidden unit activations. In both cases, the population activity was confined to lie close to a 2D manifold regardless the initial condition. Alternatively, we constructed the 3D embedding of *N* -dimensional population activity using a two-step hybrid dimensionality reduction procedure (Gardner et al. 2022): linear PCA (with the first 6 PCs) followed by a nonlinear dimensionality reduction method known as UMAP (Uniform Manifold Approximation and Projection). This visualization step revealed a 3D torus-like structure (Figure 4C for Setup #2; see Figure S5 for more results). Additionally, the ring structure did not emerge in the case of linear E/I-RNN (Figure S5F).

**Figure 4:**
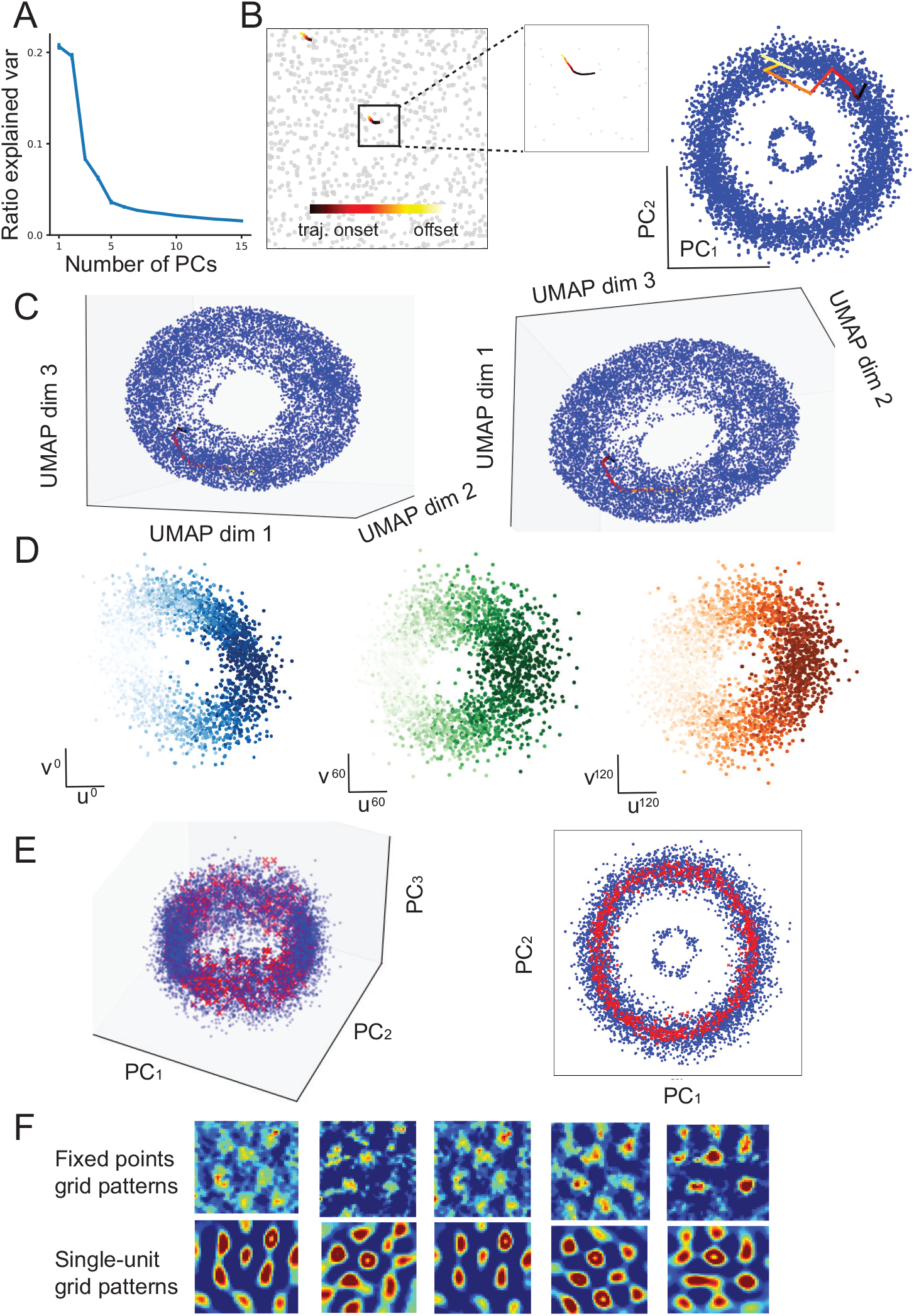
Emerged Low-Dimensional Ring Manifold and Attractor. **(A)** PCA revealed the explained variance ratio of trained RNN units. Error bar denotes the SEM from *n* = 10 trained networks. **(B)** Left: 200-ms color-coded simulated trajectory in the 2D enclosure. Right: Two-dimensional (2D) ring manifold: Projection of the high-dimensional RNN population activity onto the first two dominant PCs (PC_1_-PC_2_ plane). Each blue dot represents a temporal sample in the simulated trajectory. Neural trajectory was color coded according to the simulated trajectory in the left. **(C)** Three-dimensional (3D) torus manifold shown from two different angles. The 3D manifold was produced by PCA followed by UMAP (see STAR Methods). **(D)** Projections of the 3D manifold onto three pairs of axes (see STAR Methods). **(E)** Visualization of the identified torus-shaped attractor (red crosses represent the fixed points) in 3D (left) and 2D (right) PCA subspace. **(F)** Comparison between five fixed-point grid patterns and selected single-unit grid patterns from the trained E/I-RNN. Notice the close resemblance. Pearson’s correlation statistics between top and bottom five panels were (from left to right): 0.828, 0.781, 0.861, 0.841, 0.827. See also Figure S5 and Movie S1.

In light of low-dimensional Fourier analysis (STAR Methods), we projected the 3D manifold onto three predetermined pairs of axes (Figure 4D), each revealing the ring structure along different unit vector spaces (e.g., 0°, 60°, 120°). Similar ring structures were also found in other axis spaces (e.g., 30°, 90°, 150°, results not shown). Furthermore, we identified fixed-points of the attractor using numerical stimulations (STAR Methods). Figure 4E shows a torus attractor pattern, indicating that the dynamic system had a stable activity pattern. Therefore, the trained network was able to maintain the memory about the environment achieved by path integration. Furthermore, the fixed points also showed grid activity (Figure 4F). Interestingly, these fixed-point grid patterns resembled the activity of some hidden units in the PCA subspace. Therefore, these results suggested that the grid pattern of hidden units was controlled by these fixed-points. That is, these fixed-point patterns also generated attractor fields, and the population activity of the recurrent dynamic system converged to them and shaped the stable grid pattern.

### Robustness of Grid Patterns to Visual and Spatial Inputs

The latent network state and recurrent dynamics could be biased by the new external input. Upon the completion of E/I-RNN training, we tested how the emerged grid patterns would change with respect to the unseen visuospatial input. First, to emulate the darkness (Light OFF) condition (Setup #2), we set the visual input to zeros for the trained E/I-RNN. Surprisingly, the visual grid patterns showed very few changes (Figure 5A). This was also seen in the neural trajectories in the PCA subspace. The overall ring attractor remained similar, except for subject to a rotation (Figure 5B). Second, to emulate a new navigation environment, we switched to a new visual scene such that an identical run trajectory would have the same velocity input but different visual input in testing. Again, we observe a stable grid pattern for a wide range of visual input (Figures 5C and 5D for Setup #2; Figure S5 for Setup #5). These results suggest that the trained E/I-RNNs and underlying recurrent attractors were robust to input perturbation.

**Figure 5:**
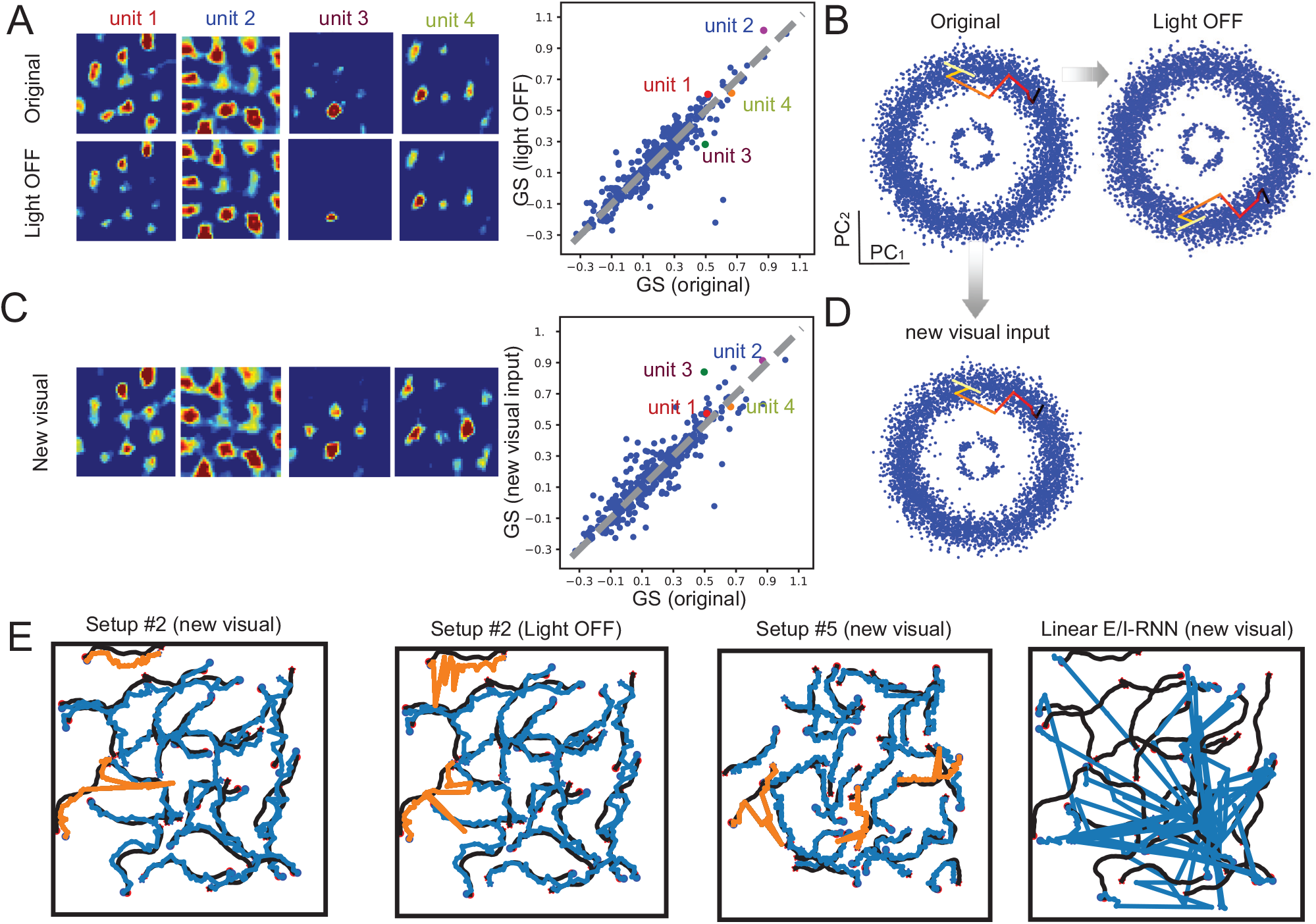
Stability of Grid Patterns and Ring Attractor. **(A)** Comparison of grid unit patterns and grid score (GS) statistics when the visual input was set to zero (light OFF) during testing (Setup #2). The GS statistics showed no statistical difference (two-sided paired signed-rank test, non-significant or n.s.). **(B)** Comparison of ring attractor and neural trajectories between the original and light OFF conditions. **(C)** Comparison of grid unit patterns and GS statistics when a new visual input was used in testing while keeping the velocity unchanged (Setup #2). The GS statistics showed no statistical difference (two-sided paired signed-rank test, n.s.). **(D)** Ring attractor and neural trajectory while using a new visual input. **(E)** In testing, the noisy long (sequence length: 50) trajectories either remained stable (blue) or first-perturbed-then-converged (orange) to the simulated paths (black) in the E/I-RNN (ReLu), suggesting the stability of ring attractor. In contrast, the noisy long trajectory tended to deviate from the simulated trajectory in the linear E/I-RNN. Snapshot examples under different testing conditions are shown: first panel: Setup #2, new visual input; second panel: Setup #2, light OFF; third panel: Setup #5, new visual input; fourth panel: Setup #2, linear activation function. The black curve denotes the simulated path, and the red circle and red star represent the start and end position, respectively. The overlaid blue curve denotes the predicted position, and the blue circle and blue star represent the start and end position, respectively. See also Figure S6.

The stability of grid patterns also predicted the stability of behavioral output in prediction. During testing, we simulated longer run trajectories (sequence length of 50) than the ones (sequence length of 10) used in the training phase. In the output space, with the same initial conditions, we found that the noisy long trajectories either remained stable or first-perturbed-then-converged to the simulated trajectories. In contrast, when the ring attractor was absent (as in the linear E/I-RNN, see Figure S5D), the trajectory output was not stable. Some snapshot examples under various testing conditions are shown in Figure 5E.

### Robustness of Grid Patterns to Recurrent Network Connectivity

In the literature, it has been suggested that the grid pattern formation can be generated through attractor dynamics in a recurrent network with a specific local connectivity including excitatory and inhibitory connections (McNaughton et al. 2006; Couey et al. 2013). The recurrent attractor dynamics of the trained E/I-RNN was primarily determined by the recurrent matrix **W**^rec^. Without additional constraints, the learned **W**^rec^ was fully connected. However, sensory cortices are known to have columnar organization: excitatory neurons within a columnar structure (also known as the hypercolumn) are densely connected and form functionally similar cortical maps (Buxhoeeveden 2002), whereas between-columnar neurons are sparsely connected. To make our E/I-RNN model more biologically realistic, we further modified the structural synaptic connectivity and sparsity (i.e., the percentage of 0 in connection weights 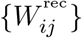).

#### Clustered E-E connectivity

The recurrent network connectivity was motivated by anatomical evidence for non-uniform or clustered connections between pyramidal neurons in the cortex (Litwin-Kumar and Doiron 2012). Previous studies have shown clustered or columnar structures in the primary and secondary areas of visual cortex (Horton and Adams 2005; Laramee et al. 2013). Therefore, we first split the E-population into two subnetworks *E*_1_ and *E*_2_ a priori, and kept such clustered connectivity during the course of RNN training (Type 1, Figures 6A top panel). Upon the completion of E/I-RNN training, we again observed robust grid responses in both E-and I-neuronal populations. Interestingly, both grid-like and band-like patterns emerged from the clustered subnetworks. This emerging continuous spectrum of spatially periodic units from the structured E/I-RNN reflect different combinations of a small set of elemental periodic bands (Krupic et al. 2012; Narvatilova et al. 2016). Additionally, the periodic grid or band-like units within the same excitatory subnetwork tended to group together (i.e.,wit similar spatial frequency or orientation); whereas inhibitory grid units tended not to group together (Figure 6B). This was conceptually consistent with the observation seen in the non-structured networks (Figures 3C and 3D). Such clustered grouping based on similar spatial receptive fields (i.e., spatial frequency and orientation) was reminiscent of the minicolumns in the visual cortex. These periodic non-localized band-like units were different from the spatially localized ON/OFF visual receptive fields.

**Figure 6:**
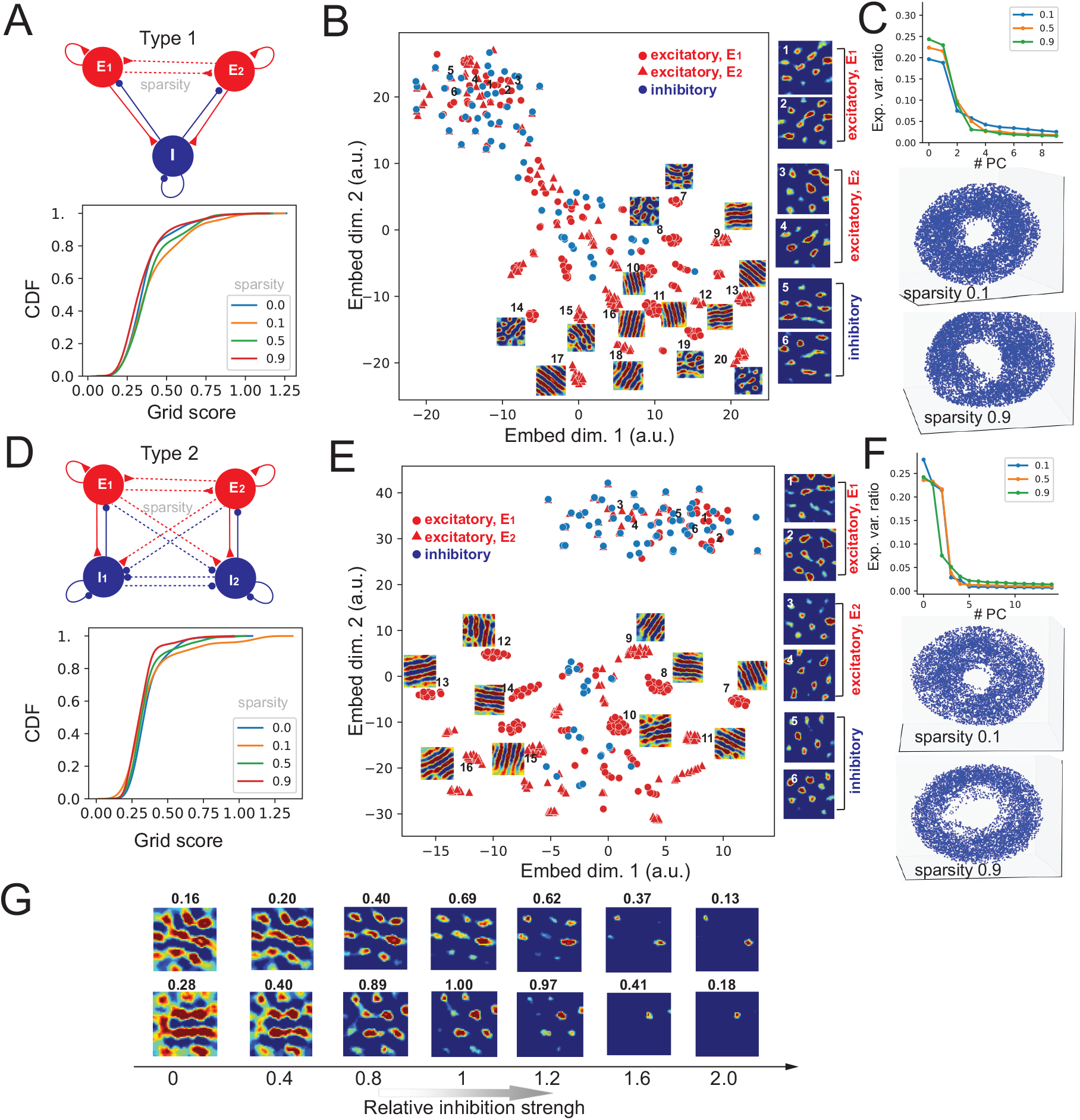
Robust Grid Patterns with Respect to Subnetwork Connectivity. **(A)** Grid score distribution statistics in Type-1 subnetwork connectivity. The dashed line denotes weak connections with various sparsity levels (0, 0.1, 0.5, 0.9). Sparsity level 0 implies full connections (original setting). Here, excitatory neurons were divided into two subnetworks: *E*_1_ and *E*_2_. **(B)** 2D projection of RNN population responses (top 50% grid scores) via the t-SNE algorithm (sparsity level: 0.5). Representative grid fields are shown from individual functional groups. **(C)** PCA explained variance ratio (top) and 3D ring manifold of RNN population responses for two sparsity levels 0.1 and 0.9 in Type-1 subnetwork. **(D)** Grid score distribution statistics in Type-2 subnetwork connectivity. Here, excitatory neurons were divided into two subnetworks: *E*_1_ and *E*_2_, and inhibitory neurons were divided into two subnetworks: *I*_1_ and *I*_2_. The E-to-I, I-to-I and I-to-E connections were weakly coupled. **(E)** Similar to panel B, except for Type-2 subnetwork connectivity (sparsity level: 0.9). **(F)** PCA explained variance ratio (top) and 3D ring manifold of RNN population responses for two sparsity levels 0.1 and 0.9 in Type-2 subnetwork. **(G)** Changes of grid-like example unit firing patterns with increasing or decreasing E/I balance. Number on the top of each grid field denotes the calculated grid score. See also Figures S7 and S8, and Movie S2.

Furthermore, in training we gradually changed the sparsity of inter-subnetwork connectivity and investigated the impact of sparsity on grid patterns. The sparsity level varied between 0 and 1, with 0 meaning fully connected (i.e., original setup), and 1 meaning completely disconnected between two subnetworks. Our results showed that the population grid score statistics remained relatively stable regardless of the sparsity level (Figures 6A, bottom panel). Additionally, the 3D ring-like manifold remained stable with increasing sparsity of inter-subnetwork connectivity (Figure 6C). Notably, an increasing higher degree of sparsity in inter-subnetwork connectivity would produce more isolated clusters within *E*_1_ and *E*_2_ groups, many of which showed band-like patterns (Figures 6B, 6E and S7).

#### Clustered I-I connectivity

Inhibitory projections were usually not clustered in our RNN model, consistent with studies showing that inhibitory neurons connect densely and non-specifically to pyramidal neurons (Litwin-Kumar and Doiron 2012). However, it has been shown that the inhibitory-to-inhibitory connections underlying a disinhibitory microcircuit may plays an important role in reshaping the recurrent dynamics (Kim and Sejnowski 2021). To test the impact of inhibitory connectivity, we further split the inhibitory population into two subnetworks *I*_1_ and *I*_2_, with each projecting to specific excitatory subpopulations. We assumed that two inhibitory subnetworks were weakly fully coupled, and but the excitatory-to-inhibitory and inhibitory-to-excitatory connections are weakly coupled (Type 2, Figures 6D top panel). Again, we varied the sparsity of inter-subnetwork connectivity and further compared their grid score statistics, grid field embedding and ring manifold structure (Figures 6D, 6E and 6F). Interestingly, we observed a mixture of band-like and grid-like patterns, yet similar band-like patterns tended to group together within the same excitatory subnetwork. Similar observations were also found for other types of structured network connectivity (see Figure S7).

#### E/I balance

The E/I balance of our trained E/I-RNN was controlled by the relative degree of excitation and inhibition. To investigate the role of inhibition in reshaping the spatial tuning of hidden units, we randomly selected excitatory grid-like units and modified their relative inhibition by gradually decreasing or increasing the inhibitory-to-excitatory input or connection strength (STAR Methods). As a result, the grid pattern and grid score changed accordingly (Figure 6G). Generally, the grid-like patterns of excitatory units had tendency to evolve into band-like patterns with decreasing inhibition strength, yet the grid patterns became weak or diminished with increasing inhibition strength. These empirical observations may explain the clustered band-like firing patterns under various structured network connectivity and sparsity levels.

#### Grid remapping

It has been well known that the firing patterns of mEC grid cells change following environmental changes, such as translation and/or rotation of fields (Hafting et al. 2005; Fyhn et al. 2007). To emulate this remapping condition, we conducted several manipulations in multiple testing conditions and examined the change of grid unit responses. First, we rotated the environment by 90 degrees, and changed all place fields accordingly by 90° rotation. Second, we proportionally increased or decreased the size of place fields in the output (to emulate a larger or smaller environment size). Third, we permuted the trained RNN output order (or equivalently, permuting the columns of **W**^out^) while keeping the place cell coverage unchanged. In the first condition, we found the grid patterns also rotated 90°, but the grid score statistics remained unchanged (Figure S8A). In the second condition, the grid patterns displayed either zoom-in or zoom-out grid patterns within the original environment, but the overall grid patterns were preserved (Figure S8B). In the third condition, we found most grid patterns changed or remapped (Figure S8C). At the population representation level, in nearly all conditions the ring manifold structures remained similar to the original one.

### Multistable Attractors Emerged from Sparsely Connected RNNs

The structural connectivity density of inter-subnetwork and intra-subnetwork determines the overall sparsity of network connectivity. Next, from a pretrained E/I RNN (with a medium sparsity of 0.5), in testing we randomly set a small percentage of inter-subnetwork or intra-subnetwork EE connection weights to zeros. This would give us an opportunity to test the “corrupt” versions of the trained RNN with disrupted prewired synaptic connectivity. Interestingly, we found that with increasing sparsity in the inter- and intra-subnetwork connectivity of the pretrained E/I-RNN, two or three isolated 3D torus structures emerged (Figures 7 and S9), although the individual grid patterns remained relatively stable. The presence of multiple-loop torus suggested a bifurcation phenomenon and the possibility of multistable attractor states (Patra 2018; Song et al. 2019). Together, these results from Figure 6 and Figure 7 suggest that functionally distinct grid patterns and multistable attractors emerged from weakly coupled excitatory subnetworks.

**Figure 7.**
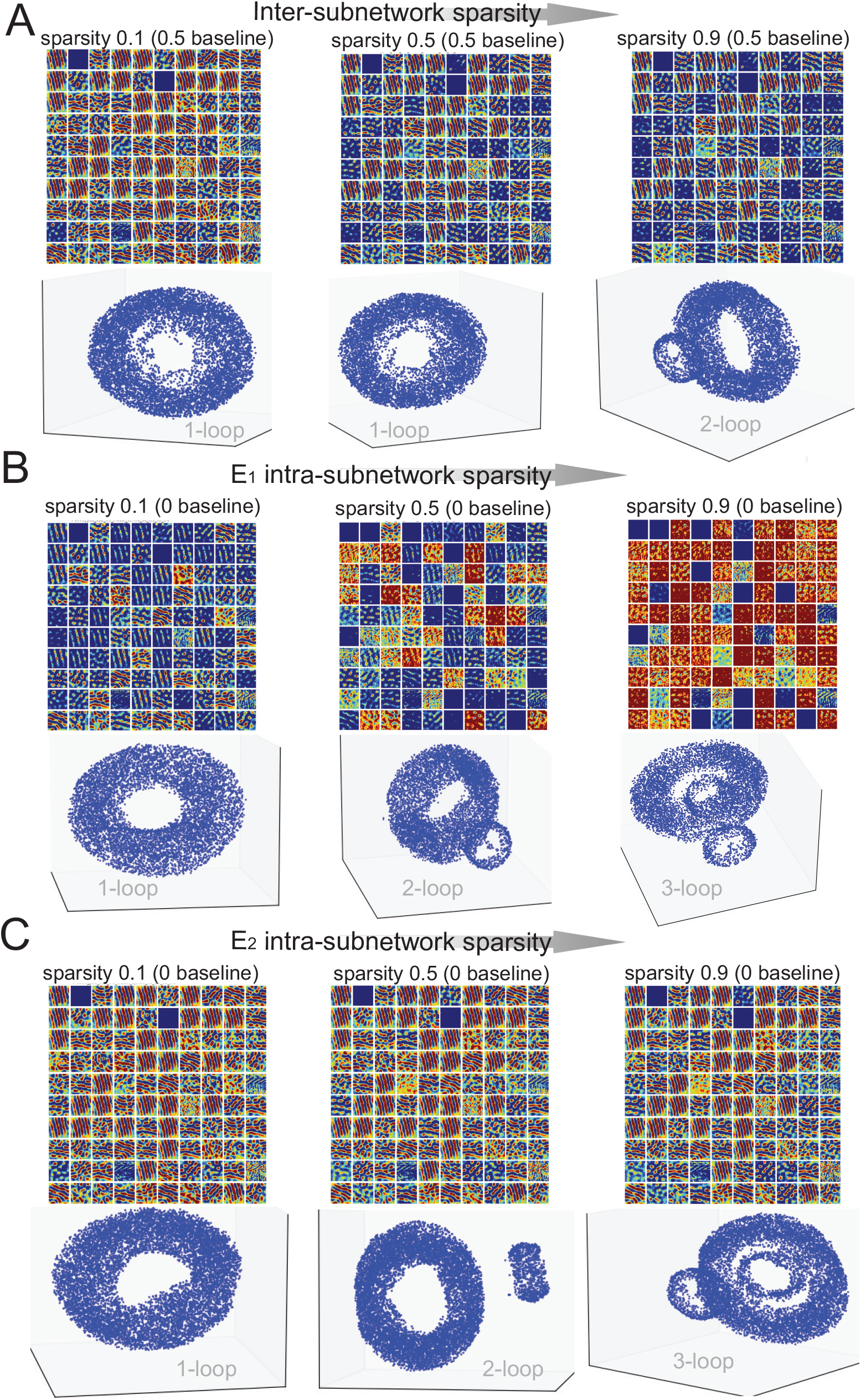
Emerged Multistable Ring Attractors with Increasing Sparsity in Network Connectivity. **(A)** Changes in 100 unit firing patterns of *E*_1_ subnetwork (top) and evolution of the 3D manifold structure (bottom) with increasing inter-subnetwork connectivity (Type 1, Setup #2, sparsity baseline 0.5). **(B)** Changes in 100 unit firing patterns of *E*_1_ subnetwork (top) and evolution of the 3D manifold structure (bottom) with increasing *E*_1_ intra-subnetwork connectivity, whereas the *E*_2_ intra-subnetwork connectivity remained unchanged (Type 1, Setup #2, sparsity baseline 0). **(C)** Changes in 100 unit firing patterns of *E*_1_ subnetwork (top) and evolution of the 3D manifold structure (bottom) with increasing *E*_2_ intra-subnetwork connectivity, whereas the *E*_1_ intra-subnetwork connectivity remained unchanged (Type 1, Setup #2, sparsity baseline 0). See also Figures S9 and S10, and Movie S3.

To get more insight into this phenomenon, we computed quantifiable statistics of learned **W**^rec^ with increasing sparsity from one trained RNN (Setup #2). We also applied Schur decomposition to the non-normal matrix **W**^rec^ and quantified the strength of functionally feedforward connections (FFCs, denoted as *κ*) as well as the eigenvalue statistics that characterize short-term and long-term behaviors (STAR Methods and Supplemental Information). We found that the maximum eigenvalue of 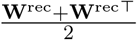 that characterizes the short-term behavior monotonically decreased with gradually increasing sparsity in both inter-subnetwork and intra-subnetwork connectivity (Figure S10A). In contrast, the maximum (real-part) eigenvalue of **W**^rec^ that characterizes the long-term behavior decreased with gradually increasing sparsity in inter-subnetwork but not intra-subnetwork connectivity. Additionally, *κ* reached a maximum with an intermediate sparsity level of inter-subnetwork or intra-subnetwork connectivity (Figure S10B). Since the largest (real-part) eigenvalue was greater than 1, it suggested that the chaotic state or bifurcation might be present in the RNN, which might explain the phase transition of multiple loops in the torus manifold.

### RNN Grid-Like Patterns Emerged in a Non-spatial Task

The next question arises whether grid patterns may emerge from the RNN that performs a non-spatial task? To investigate this question we considered a modified MNIST handwritten digit sequence recognition task. We first embedded the high-dimensional visual stimuli into a 2D space in the output. In the output space, the images were grouped together in the embedded space according to their similarity. Navigating between points in the embedded space can be viewed as a sequence recognition task. We employed a pretrained convolutional neural network (CNN) that emulated early-stage V1 preprocessing (Figure 8A), and subsequently fed the flattened features of the CNN as input to the E/I-RNN (STAR Methods). Only the RNN parameters were trained in the CNN-RNN architecture to map the image sequence to the 2D coordinate in the embedded space. We envisioned that when a set of digital image sequences (each with sequence length 10) was presented to the input of the CNN-RNN, the output produced a trajectory in the embedded space. Two differences arose from the previous computer simulation experiments: first, the embedding coordinate space was not uniformly sampled by the trajectories; second, the step size of trajectory was non-even, with random speed and direction at every single step.

**Figure 8:**
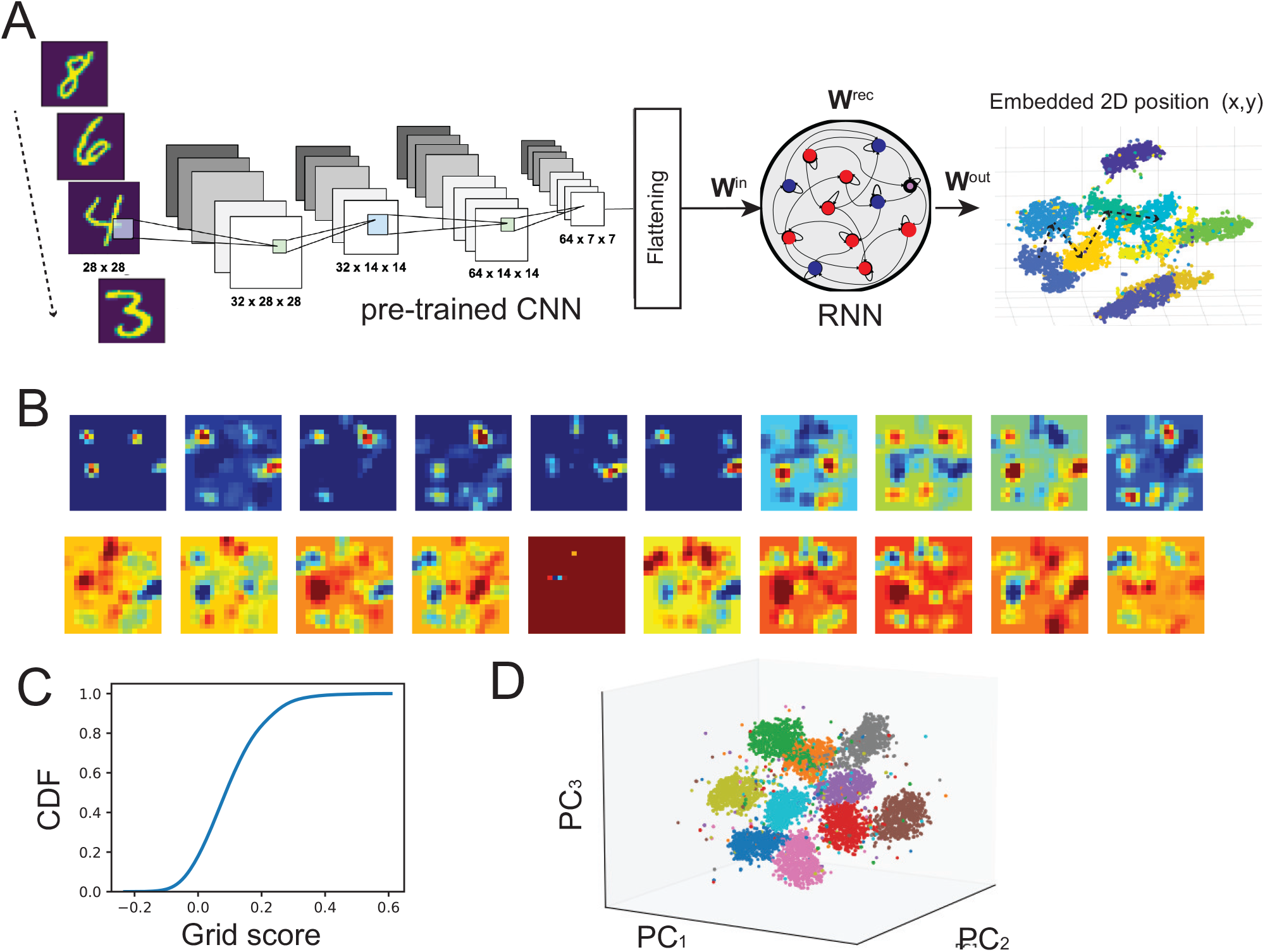
Visual Grid Patterns Emerged in an CNN-RNN Model for Non-spatial Task. **(A)** Schematic of the proposed CNN-RNN model to perform a visual recognition task. The CNN model was pretrained and the flattened features was fed to the RNN. **(B)** Examples of visuospatially-modulated units in the trained RNN. **(C)** Cumulative distribution function (cdf) of the grid score from the trained E/I-RNN hidden units. **(D)** Low-dimensional manifold in the PCA subspace, with each color representing one digit.

Upon the completion of training the RNN, we repeated the same analysis as in the spatial task. At the single unit level, we found a wide range of spatially tuned units. Some displayed non-periodic irregular grid-like patterns, and others showed random patterns (Figure 8B). As a result, the grid score statistics of the whole population was lower (Figure 8C). In this case, the low-dimensional population response formed a 10-cluster manifold structure in the PCA subspace (Figure 8D).

## DISCUSSION

Numerous evidence and publications have recently pointed to rich spatial modulation across a wide range of visual thalamus and cortices (Hok et al. 2018; Ji and Wilson 2007; Haggerty and Ji 2015; Saleem et al. 2018; Fournier et al. 2020; Campbell and Giocomo 2018; Flossmann and Rochefort 2021; Long et al. 2021a; Zong et al. 2022). A preliminary finding has suggested a compact spatial map that consists of place cells, gird cells, head-direction cells, and border cells exists in the rat V2 visual cortex (Long et al. 2021a). However, the sources and functional role of these spatially-modulated signals in primary and secondary visual cortices remain a puzzle. Theta oscillations and theta-modulated firing have been found in the rat V2 (Long et al. 2021a), providing a possible driving force of speed input. Additionally, anatomy projection between the V2 and V1, and projections from the secondary motor (M2) and retrosplenial (RSC) cortices may provide additional self-motion, place-modulating and directional inputs to visual cortices. Furthermore, spatial modulation of place cells and grid cells in the rat V2 persisted in the absence of sensory input (e.g., darkness), suggesting the robustness of these spatially modulated neurons. These results were consistent with the prior experimental findings in rodent mEC grid cells (Hafting et al. 2005; Barry et al. 2012; Dannenberg et al. 2020).

To date, the receptive fields of V2 neurons have been studied using only pure visual stimuli. For instance, V2 neurons in monkeys could be broadly classified as V1-like (typical Gabor-shaped subunits), with ultralong (subunits with high aspect ratios) or complex-shaped (subunits with multiple oriented components) (Liu et al. 2016). There is also evidence that several rodent extrastriate areas process information related to other sensory modalities (Miller and Vogt 1984; Sanderson et al., 1991). This may indicate fewer hierarchical stages in the rodent, relating visual information more readily to multimodal interactions or complex behaviors such as spatial navigation (Marshell et al. 2011). Without directly modeling visual receptive fields, we found independence between spatial tuning and visual tuning among the emerged grid units in the trained RNNs, suggesting the possible task-dependent mixed selectivity in generalized grid codes (beyond mEC or V2 grid cells) during complex task behaviors (Klukas et al. 2020). Structural network connectivity is also linked to functional clustering. In the V1, it has been known that neurons with similar nonclassical extra-receptive fields (ERFs) have tendency to group into clusters (Yao and Li 2002); these clusters are randomly distributed in all cortical layers, with no detectable relationship with orientation and ocular dominance columns. Our results that clustered grid/band-like patterns emerge from sparse structural RNN connectivity support this intuition. However, experimental verification of this model prediction would require high-density large-scale recordings from the rat V2.

In our biologically-constrained RNN model, we envisioned that the output layer that encodes the place is the V1, which serves as a teaching signal to the V2. This is not impossible because first, the rodent V1 neurons are known to have place tunings (Ji and Wilson 2007; Haggerty and Ji 2015; Fournier et al. 2020). Second, the back-propagating error from the output V1 units to the recurrent V2 units represents the information flow in the visual pathway. Third, the V2 also provides a modulation input to the V1 in the feedback pathway (De Pasquale et al. 2013). Our RNN-based V2 grid cell model distinguishes itself from other mEC grid cell models in several noticeable ways. First, we generalized the standard RNN models (Cueva and Wei 2018; Sorscher et al. 2020) by incorporating Dale’s principle and structured intra-cortical connectivity. These constraints make our model more biologically realistic in relation to the V2 visual cortex. Second, we showed that the visual optical flow can be used an alternative input to the V2 as an analogue of the velocity input, which both yielded robust grid patterns and torus-like attractors. The optical flow can be implemented in the early stage of visual system (Wang et al. 1989; Wurtz 1998), and the heading direction can be estimated from optical flow in the visual cortex (Lappe and Rauschecker 1992) or sensorimotor circuit (Leinweber et al. 2017). Furthermore, our E/I-RNN models reveal torus-like manifolds and attractors in a low-dimensional space, consistent with the other experimental and computational findings (Gardner et al. 2022; Sorscher et al. 2020). Interestingly, multistable torus-like manifolds may emerge from the E/I-RNN with increasing inter- or intra-subnetwork connectivity, suggesting that sparsely connected biological networks may use multiple ring attractors to store information.

Predictive maps have been suggested as the common computational principles for generating “place codes” and “grid codes” in several computational models (Stachenfeld et al. 2017; Recanatesi et al. 2021). Specifically, the successor representation (SR) (Dayan 1993) is the product of the inherent state-action transition dynamics that characterizes a predictive dynamics. In the context of spatial navigation, the SR for a given state (a spatial location) is radially symmetric over space, and the columns or eigenvectors of the SR matrix correspond to the place fields and grid fields, respectively (Stachenfeld et al. 2017; Gershman 2018). The state-action transition matrix bears a functional resemblance with the recurrent weight matrix in the RNN that characterizes the network state transition. Our work is also in spirit close to the recent RNN modeling effort (Recanatesi et al. 2021), in which the RNN was trained to predict the future action based on the current state and action, and the trained recurrent weight matrix that characterizes a predictive representation showed emergent place-like response patterns. Unlike other continuous attractor networks or continuous ring attractor models, the network connectivity matrix of our E/I-RNN is asymmetric and non-normal. Our results have shown that such a recurrent network structure produces robust grid patterns and torus-like attractors under various input configurations and network connectivity. Recently, the RNN has been demonstrated to be capable of constructing a ring manifold that constrains a set of discrete fixed points (Pollock and Jazayeri 2020). We speculate that such synthesis-based approach may potentially be useful for modeling biomodal attractor dynamics in visual grid patterns (Long et al. 2021b).

Path integration relies on an egocentric coding process, which allows animals to integrate information (e.g., speed of movement, travel time, and directional change) generated by the self-movements, to update the position in relation to a start location. Visual landmarks and motion cues are also critical for visuospatial integration. In virtual reality experiments, internal vestibular (inertial) and proprioceptive cues are disrupted; however, some idiothetic cues that are not internally generated can be still used for path integration. Such cues may include visual optic flow, airstream detection (e.g. by a rat’s whiskers) or other sensory reafference inputs produced by locomotion (Savelli and Knierim 2019). An outstanding question is whether path-integration-like computations are also used in non-spatial domains, such as building non-spatial representations such as time intervals or trajectories defined in a sensory stimulus space? Multiple lines of evidence have suggested a confirmative answer. For instance, it has been shown that self-organized domain-general learning algorithms may explain the emergence of grid cells in both spatial and conceptual domains (Mok and Love 2019), and grid representations may provide efficient similarity search strategies in the generalized and continuous cognitive space. Additionally, a computational model of visual grid cells has been proposed for visual recognition memory, where a sequence of memorguided saccades can encode salient stimuli (Bicanski and Burgess 2019). Our demonstration of visual grid patterns produced from a CNN-RNN model suggests that the computational principles of grid computation is beyond the velocity-driven path integration task. The RNN has provided a dynamical systems viewpoint for motor movement (Sussilo et al. 2015; Michaels et al. 2016), path integration (Cueva and Wei 2018; Sorscher et al. 2020), information integration (Zhang et al. 2021), and predictive representations (Recanatesi et al. 2021). We envision that the recurrent dynamics of RNNs can be biologically implemented by a neural substrate in a wide range of cortical networks outside the traditional hippocampus-entorhinal system. The pursuit of grid codes in the context of perception and cognition tasks has just begun (Yu et al. 2021).

Several limitations are noticed in our biologically-inspired RNN models for visual grid patterns. For instance, our simulated visual input (either illumination or optical flow) to the RNN is rather simplified and can not fully capture the natural vision. However, we speculate that the change in visual simulation setup by adding an additional level of complexity will not affect the basic finding, since the grid responses are rather robust to different visual input configurations. Furthermore, as a stand-alone model, the E/I-RNN does not explicitly consider the cortico-cortical input in the biological brain. Despite these limitations, our computational modeling work may produce experimentally testable predictions. First, our computer simulation results suggest that the intra-cortical connectivity and E/I balance have an impact on the grid responses. This can be experimentally tested using intra-cortical microstimulation (Averna et al. 2020; Averna et al. 2021) or optogenetic stimulation (Bridi et al. 2020). Third, it remains unknown how visual grid patterns will change upon modifying the upstream input of the V2 (such as the V1 and visual thalamus). This can also be experimentally investigated by selective inactivation of these feedforward connections. Fourth, the mixed selectivity of V2 neurons can be tested via a series of experiments with controlled spatial and visual stimuli. Finally, it would be alluring to find grid patterns in visual and other sensory cortices during non-spatial tasks in rodent experiments.

## STAR Methods

Detailed methods include the following:

- KEY RESOURCE TABLE
- RESOURCE AVAILABILITY
  - Lead contact
  - Materials availability
  - Data and code availability
- EXPERIMENTAL MODEL AND SUBJECT DETAILS
  - MNIST dataset
- METHOD DETAILS
  - Input stimuli and output coding for a spatial navigation task
  - RNN structure and training for a spatial navigation task
  - CNN-RNN architecture for a non-spatial task
  - Analysis of ring manifold and attractor
- QUANTIFICATION AND STATISTICAL ANALYSIS
  - Computation of grid score, grid field and autocorrelogram
  - Computational of optical flow
  - Dimensionality reduction
  - Schur decomposition

### KEY RESOURCE TABLE

| REAGENT or RESOURCE | SOURCE | IDENTIFIER |
| --- | --- | --- |
| **Experimental Data** |  |  |
| MNIST dataset | Open source | http://yann.lecun.com/exdb/mnist/ |
| **Software and Algorithm** |  |  |
| Grid score computation | (Banino et al. 2018) | https://github.com/deepmind/grid-cells |
| t-SNE | (van der Maaten and Hinton 2008) | https://lvdmaaten.github.io/tsne/ |
| UMAP | (McInnes and Healy 2018) | https://github.com/lmcinnes/umap |
| CNN | Open source | https://github.com/iamkrut/MNIST_handwriting_classification |
| E/I RNN | (Song et al. 2016) | https://github.com/frsong/pycog |
| fixed-point finder | (Sussillo and Barak 2013) | https://github.com/mattgolub/fixed-point-finder |

### RESOURCE AVAILABILITY

#### Lead contact

Further information and requests for data should be directed to and will be fulfilled by the Lead Contact, Zhe S. Chen.

#### Materials availability

All materials are contained in the paper or available from the Lead Contact.

#### Data and code availability

Data and code are available upon request to the Lead Contact. Some custom Python scripts are available at https://github.com/Xhan-Zhang/grid-rnn/.

### EXPERIMENTAL MATERIALS AND DATA

#### MNIST dataset

The MNIST (Modified National Institute of Standards and Technology) dataset consists of 70000 28*×*28 pixel grayscale images of handwritten single digits between 0 and 9. The digits have been size-normalized and centered in a fixed-size image. It is publicly available.

#### Rat V2 experimental recordings

The experimental recordings of V2 grid cells from freely behaving rats were described in details elsewhere (Long et al. 2021a). We adapted our computer simulation setup to match animal’s behavioral statistics (such as run speed and direction). The rat local field potential (LFP) theta (8-10 Hz) rhythms from the V2 visual cortex were also used to guide the computer simulation (Setup #4). Specifically, we used the animal’s real speed and interpolate the LFP theta frequency based on a 3-rd polynomial mapping. The resultant theta frequency was ranged between 8 Hz and 9 Hz (Figure 1G).

### METHOD DETAILS

#### Input stimuli and output encoding for a spatial navigation task

In computer simulations, we generated 5000 random trajectories within the two-dimensional (2D) environment (2.2 m*×*2.2 m) to simulate the animal’s trajectories based on random speed and head direction. Each trajectory had a fixed sequence length (*ℓ* = 5-30, equivalent to 100-600 ms). The initial speed and head direction were randomized. To encode a 2D position, we assumed that the encoding of individual place unit had a 2D isotropic Gaussian shape as defined below (Banino et al. 2018):

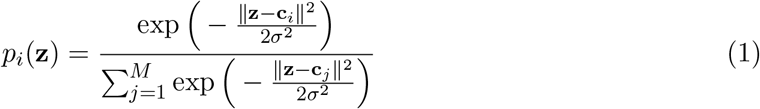

where **c**_*i*_ ∈ ℝ^2^ denotes the center of place receptive field, and *σ* > 0 denotes the place cell scale. We used *M* = 1024 units to uniformly cover the 2D environment (Fig. 2B). From the population activity of these place-modulating units, we could recover the 2D position **z** ≡ (*x, y*). Head-direction activations were sampled from a Gaussian distribution with zero mean and variance of 11.5 radians.

For Setup #1, the RNN only received the 2D velocity input at each time bin. For Setup #2, we added the additional visual input illuminance. Based on the heading direction, we defined a viewing region of interest (ROI) defined by an image of 8 *×* 8 pixels (i.e., dimensionality 64), which corresponded to the raw visual input (such as the luminance of pixels). To reduce the dimensionality, we further applied principal component analysis (PCA) and projected the vectorized image onto the dominant PC subspace. In our experiment, we tried varying number of PCs (2-20) that explained up to 93.4% variance in the visual stimuli. For Setup #3, the RNN received the 2D acceleration and visual input illuminance at each time bin. For Setup #4, we used the actual animal’s run speed in the 2D environment to generate the trajectories, and further simulated the theta frequency based on a previously reported relationship between run speed and theta frequency (Chen et al. 2016).

#### RNN structure and training for a spatial navigation task

We trained an excitatory-inhibitory (E/I) RNN to perform a simulated spatial navigation task in the 2D environmental enclosure (Fig. 2A). We assumed the *N* -dimensional neural state dynamics, **x**(*t*), was driven by a recurrent dynamics plus an *N*_in_-dimensional input **u**(*t*):

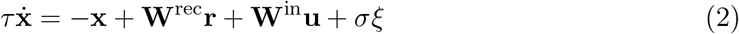

where *τ* denotes the time constant, *ξ* denotes additive *N* -dimensional Gaussian noise, each independently drawn from a standard normal distribution, *σ*^2^ defines the scale of the noise variance; **W**^rec^ is an *N × N* matrix of recurrent connection weights, and 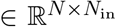 denotes the matrix of connection weights from the inputs to network units. The network produced an *N*_out_-dimensional output **z**(*t*) = **W**^out^**r**, where the neuronal firing rate vector **r** is defined by an activation function *ϕ*(**x**), which by default is a nonnegative rectified linear unit (ReLu)

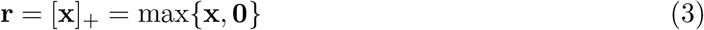

The ReLu is scale-invariant and favors sparse activation. It is noted that in a special case when the activation function is a linear unit, the firing rate dynamics will be characterized by a linear dynamical system (Rajakumar et al. 2021)

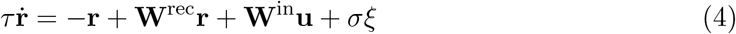

The E/I RNN was designed to satisfy the Dale’s rule such that the ratio of excitatory to inhibitory units was 4:1 (Song et al. 2016). We used *N* =512 and *dt* = 20 ms in all numerical stimulations. Depending on specific assumed input configurations (Table 1), the RNN input had different dimensionality, and the RNN output consisted of *N*_out_=1024 place-modulated units, whose linear readout produced the 2D spatial position.

The recurrent weight matrix consisted of four functional submatrices according to the cell types: excitatory-to-excitatory (EE), inhibitory-to-excitatory (IE), excitatory-to-inhibitory (EI), and inhibitory-to-inhibitory (II) connections. Generally, **W**^rec^ is non-normal (unless all submatrices {**W**_EE_, **W**_EI_, **W**_IE_, **W**_II_} are symmetric and EI and IE connections are identical); as a result, its eigenvectors are not mutually orthogonal (Goldman 2009; Murphy and Miller 2009). For individual postsynaptic excitatory or inhibitory unit, the net excitatory and inhibitory currents was summed by the presynaptic input as follows

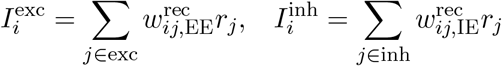

where *r*_*j*_ = [*x*_*j*_]_+_ = max(*x*_*j*_, 0) denotes the neuronal firing rate of the *j*-th presynaptic neuron, 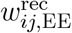 and 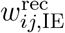 represent the EE and IE weights within **W**^rec^, respectively.

We employed the mean squared error as the cost function. The RNN was trained by back-propagation through time (BPTT) using the Adam algorithm with default configuration of hyperparameters. In batch training, we used a learning rate of 0.0005 and a batch size of 256 randomly generated run trajectories. In each input configuration, we trained at least 10 RNNs with independent initializations in parallel via GPU. In the paper, we reported the representative results from one or ten trained networks.

#### CNN-RNN architecture for a non-spatial task

In the non-spatial task, we applied stochastic neighbor embedding (SNE) to the MINIST handwritten digit images and projected the images of 28 *×* 28 pixels onto a low-dimensional (2D or 3D) space. We used the t-distributed SNE algorithm and color coded different classes of digits in visualization (van der Maaten and Hinton 2008).

We used a pretrained convolutional neural network (CNN) to emulate early visual processing in the V1 visual cortex. The simple CNN architecture consisted of two convolution layers, each followed by max pooling operations. Each unit used a ReLu activation function. In our setting, we discarded fully connected layer in the pretrained CNN (for classification); instead, we used the flattened input and fed that into the E/I-RNN to perform a visual navigation task in the embedded space (Fig. 8A). In the new task, the pretrained CNN parameters were fixed, and the E/I-RNN parameters were modified using a similar optimization procedure as in the spatial navigation task. The visual sequence consisted of 10 randomly permuted handwritten digit images (0-9) in the feature space; a total of 500,000 sequences were used to train the E/I-RNN.

#### Analysis of ring manifold and attractor

In order to identify the low-dimensional structure of population activity in the trained RNN, let **r**_*i*_(*x, y*) denote the *i*-th unit firing rate map with respect to the 2D location (*x, y*); we computed its 2D (spatial) Fourier transform

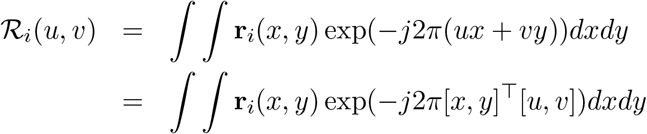

where (*u, v*) denote the spatial frequencies. The function *ℛ*_*i*_(*u, v*) is constant when [*x, y*]^⊤^[*u, v*] = (*ux* + *vy*) is constant; the magnitude of the vector (*u, v*) produces a frequency, and its phase gives an orientation. The function is a sinusoid with this frequency along the direction, and constant perpendicular to the orientation. The maxima and minima of real-valued sinusoidal basis cos 2*π*(*ux* + *vy*) occur when the inner product 2*π*[*x, y*]^⊤^[*u, v*] = *nπ* corresponds to a set of equally spaced parallel lines, which have wave-length 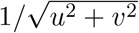 and are perpendicular to vector [*u, v*].

We further computed three spatial phases for each unit’s rate map as follows (Sorscher et al. 2020)

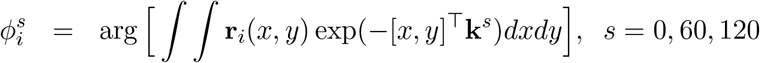

where **k**^0^, **k**^60^, **k**^120^ represent the 0°, 60° and 120° rotation unit vectors, respectively. The inner product between [*x, y*] and the rotation vector yields a new set of coordinate (*x*_*new*_, *y*_*new*_):

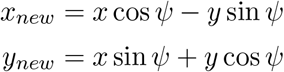

where *ψ* denotes the rotation angle. Finally, we projected the population activity onto the three orthogonal pairs of axes: *{u*^0^ ≡ cos(*ϕ*^0^), *v*^0^ ≡ sin(*ϕ*^0^)*}, {u*^60^ ≡ cos(*ϕ*^60^), *v*^60^ ≡ sin(*ϕ*^60^)} and *{u*^120^ ≡ cos(*ϕ*^120^), *v*^120^ ≡ sin(*ϕ*^120^)}. Note that the choice of (0°, 60°, 120°) was based on the assumption of a perfect hexagonal grid; for the quadrilateral grid, the choice would be (0°, 90°, 180°). In practice, we found that a wide range of rotation angles gave qualitative similar results.

To identify the attractor or slow point of the low-dimensional dynamics, let 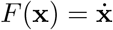 in equation (2); the Jacobian of the RNN was computed as

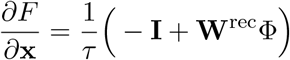

where Φ represents an *N* -by-*N* diagonal matrix containing the derivative of *ϕ*(**x**) with respect to **x**. Applying eigenvalue decomposition to the Jacobian matrix, we obtained *N* eigenmodes (eigenvectors) in the state space, and the associated *N* complex-valued eigenvalues that quantified the rate and direction along individual dimension (Pollock and Jazayeri 2020). At the fixed points, we set *F* (**x**^∗^) = 0, or equivalently

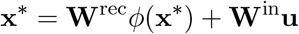

The analytic solution is not available because of the ReLu nonlinearity *ϕ*(**x**). In the special case of linear RNN, the fixed point was given by **x**^∗^ = (**I** − **W**)^−1^**W**^in^**u**. When (**I** − **W**) has a full rank, then the fixed point is unique; otherwise, the linear RNN has more than one fixed point (Seung 1998). In our analysis, we identified fixed-points or slow points by numerically solving the optimization problem (Sussillo and Barak 2013):

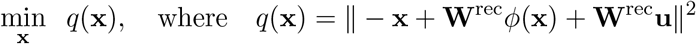

We collected a set of fixed-points by randomly initializing the network on a grid of 100 *×* 100 spatial locations in the 2D environment. Specifically, the dimensionality of the fixed point was the same as *dim*(**x**); once the numerical optimization was completed, we applied PCA to visualize the fixed points in the three- or two-dimensional PC subspace.

### QUANTIFICATION AND STATISTICAL ANALYSIS

#### Computation of grid score, grid field and autocorrelogram

The grid score (GS) was previously established to quantify the grid cell (Hafting et al. 2005; Banino et al. 2018). To be consistent with all setup conditions, we set an empirical GS threshold of ≥0.3 for the grid unit. The empirical threshold was guided by on a random shuffle distribution using a Monte Carlo *P <* 0.01. If a unit was categorized as a grid cell, the grid field was defined as the spatial rate map (50 × 50 bins) normalized by the behavioral occupancy. Using a similar method (Banino et al. 2018), we calculated the spatial autocorrelation with smoothed rate maps. Let *r*(*x, y*) denote the unit’s mean firing rate at a two-dimensional Cartesian coordinate (*x, y*), the autocorrelation of the spatial firing field was calculated as (Long and Zhang 2021):

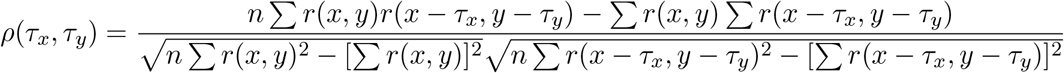

where the summation was over *n* pixels for both *r*(*x, y*) and *r*(*x* − *τ*_*x*_, *y* − *τ*_*y*_) (where *τ*_*x*_ and *τ*_*y*_ denote the spatial lags).

#### Computation of optical flow

Optical flow is commonly referred to the pattern of apparent motion of objects in a visual scene. In computer vision, the optical flow methods try to calculate the motion between two image frames which are taken at times *t* and *t* + Δ*t*. Based on local Taylor series approximations of the frame images, these methods use partial derivatives with respect to the spatial and temporal coordinates. For a 2D+*t* dimensional case, if a voxel at location (*x, y, t*) with visual illuminance *I*(*x, y, t*) are moved by (Δ*x*, Δ*y*, Δ*t*) between two image frames, then the following “brightness constancy constraint” needs to be satisfied

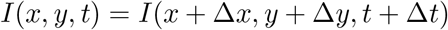

Based on first-order Taylor series expansion, the following equation can be derived

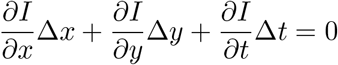

or

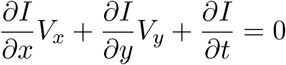

where 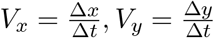. We then used the Horn-Schunck estimation method to estimate the optical flow based on neighboring frames (Horn and Schunck 1981; Cao et al. 2021). The optical flow was represented and visualized by a 2D vector field, with arrows indicating the direction, and the size of arrow proportional to the scale. In our experiment, we used a 16 × 16 visual frame to compute the optical flow, resulting in a 512-dimensional vectorized feature.

#### Dimensionality reduction

To visualize the low-dimensional recurrent dynamic attractor, we applied linear-based principal component analysis (PCA) and projected the *N* -dimensional latent vector onto a 2D subspace (PC_1_-PC_2_ or PC_2_-PC_3_) according to the percentage of explained variance. In addition, we used a nonlinear-based dimensionality reduction technique known as UMAP (Uniform Manifold Approximation and Projection) (McInnes and Healy 2018; Becht et al. 2019). The algorithm is designed to find an embedding by searching for a low dimensional projection of the data that has the closest possible equivalent fuzzy topological structure.

To quantify the similarity of grid fields, we embedded the 50 × 50 grid maps into a 2D space using the t-SNE algorithm (van der Maaten and Hinton 2008), with a default perplexity parameter of 30.

#### Schur decomposition

We applied Schur decomposition to the trained weight connectivity matrix: **W**^rec^ = **QTQ**^−1^, where **Q** is a unitary matrix whose columns contain the orthogonal Schur mode, and **T** is a lower triangular matrix that contains the eigenvalues along the diagonal (Goldman 2009; Murphy and Miller 2009; Rajakumar et al. 2021). The triangular structure of **T** can be interpreted as transforming a RNN into a feedforward neural network, and the recurrent matrix **W**^rec^ corresponds to a rotated version of the effective feedforward matrix **T**, which defines self-connections and functionally feedforward connections (FFCs) of the neural network. Unlike eigenvalue decomposition, the Schur decomposition produces the simplest (yet non-unique) orthonormal basis for a non-normal matrix. The Schur decomposition of the non-normal matrix **W**^rec^ naturally provides a separation of “diagonal” (recurrent) and “non-diagonal” (feedforward) parts. To quantify the strength of FFCs (denoted by *κ*), we computed the sum of absolute squares of the off-diagonal elements of **T**, and normalized by the sum of absolute squares of all the elements of **T** (Murphy and Miller 2009):

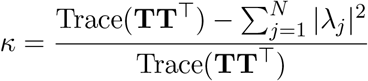

where 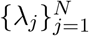 denote the eigenvalues of the lower diagonal matrix **T**. The value *κ* is interpreted as the proportion of dynamics driven by FFCs, whereas 1 − *κ* is interpreted as the proportion of dynamics driven by functionally recurrent connections.

## SUPPLEMENTAL INFORMATION

Supplemental information will be provided online.

## ACKNOWLEDGMENTS

Z.S.C. and X.Z. were partly supported by the US National Institutes of Health (R01-MH118928 and R01-NS121776). Z.S.C. also received cloud computing resources supported by the Oracle for Research Award.

## AUTHOR CONTRIBUTION

Z.S.C. conceived the study; Z.S.C. and X.Z. designed the experiment; X.L. and X.-J.Z. provided animal data recordings that motivated the computational modeling study; X.Z. performed all computer experiments and analyses; Z.S.C. wrote the initial draft of the manuscript; Z.S.C., S-J. Z. and X.Z. edited and reviewed the final manuscript; Z.S.C. acquired the funding; Z.S.C. supervised the project.

## SUPPLEMENTAL INFORMATION

### Linear RNN as a path integrator

Let’s consider a continuous-time vector differential equation that describes the dynamics of the linear RNN with *N* recurrent connected neurons:

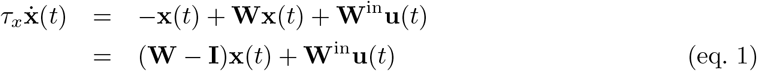

For simplicity, let the time constant *τ*_*x*_ = 1 and set **A** = **W** − **I** (where **I** is an *N × N* identity matrix), then

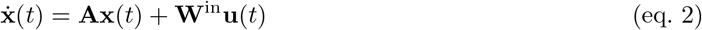

Given the linear dynamical system, we define a matrix function Φ(*t, τ*) that has the following two properties

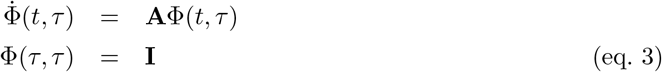

where the matrix function is referred to as the state transition matrix. If **A** is non-singular, then the state transition matrix is unique. Solving the linear vector differential equation yields

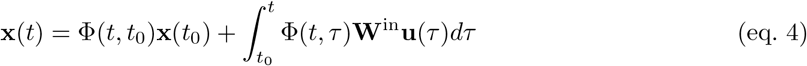

When the input **u**(*t*) is completely absent, the dynamical system is purely driven by the recurrent dynamics governed by the eigenfunctions of **A**:

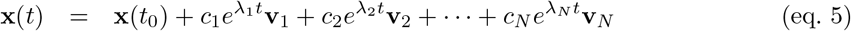

where {*λ*_1_, …, *λ*_*N*_ } are the eigenvalues, and {**v**_1_, …, **v**_*N*_ } are the eigenvectors of the matrix **A**.

The linear readout of the RNN output is given as

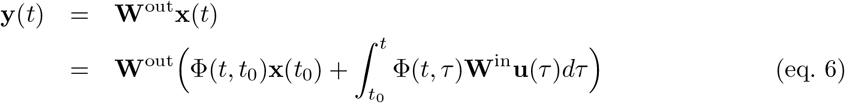

### Non-normal Connectivity and Dynamics

We call a matrix **A** *normal* if it it satisfies **AA**^⊤^ = **A**^⊤^**A**, and a stable linear normal system is contractive (Murphy and Miller 2009). On the other hand, a matrix **A** is non-normal if it satisfies **AA**^⊤^ ≠ **A**^⊤^**A**. The recurrent weight matrix **W**^rec^ of the E/I-RNN is asymmetric and non-normal, unless all submatrices {**W**_EE_, **W**_EI_, **W**_IE_, **W**_II_} are symmetric and the EI and IE connections are identical. As a result, the eigenvectors of **W**^rec^ do not form the orthonormal bases (Murphy and Miller 2009).

Specifically, an arbitrary recurrent weight matrix **W**^rec^ can be rewritten in the following form

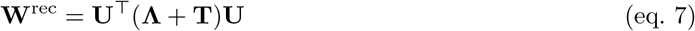

where **U** = *{u*_*ij*_} is unitary, and **Λ** is a diagonal matrix that contains the eigenvalues *{λ*_*k*_} of **W**^rec^, and **T** is a lower-diagonal matrix. The vectors of **U** are called the Schur vectors (or Schur modes) and are mutually orthogonal. In the linear E/I-RNN, let 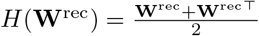 be the Hermittian part of the recurrent weight matrix, then the maximum of the eigenvalue of *H*(**W**^rec^) characterizes the short-term behavior of the network undergoing a transient growth before asymptotically converging to zeros (Asllani et al. 2018). Additionally, the maximum of the real (or real-part) eigenvalues of **W**^rec^ characterizes the long-term behavior for the speed of network’s steady state decaying to zero. Highly non-symmetric interactions of simulated neurons may create non-normal dynamics, such as large transients (Kerg et al. 2019).

**Figure S1:**
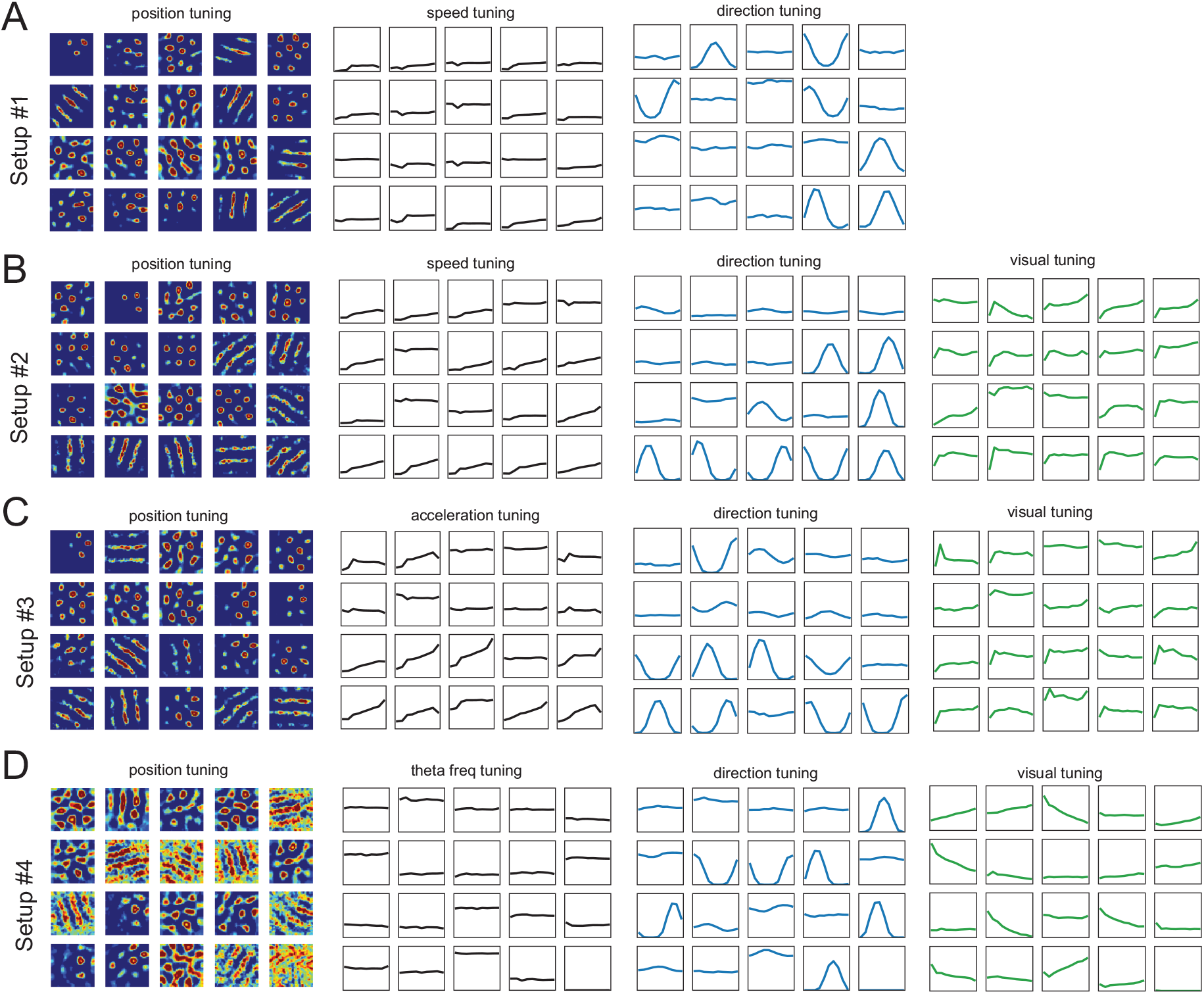
Examples of heterogeneous responses of RNN units; related to Figure 1. **(A)** Heterogeneous unit tunings from a trained E/I-RNN (setup #1). The position-modulated, speed-modulated, and direction-modulated are shown from left to right, respectively. All tuning curves are displayed with the same vertical scale (a.u.). **(B)** Heterogeneous unit tunings from a trained E/I-RNN (setup #2). The position-modulated, speed-modulated, direction-modulated, and visual illumination-based tunings are shown from left to right, respectively. **(C)** Heterogeneous unit tunings from a trained E/I-RNN (setup #3). The position-modulated, acceleration-modulated, direction-modulated, and visual illumination-based tunings are shown from left to right, respectively. **(D)** Heterogeneous unit tunings from a trained E/I-RNN (setup #4). The position-modulated, theta frequency-modulated, direction-modulated, and visual illumination-based tunings are shown from left to right, respectively.

**Figure S2:**
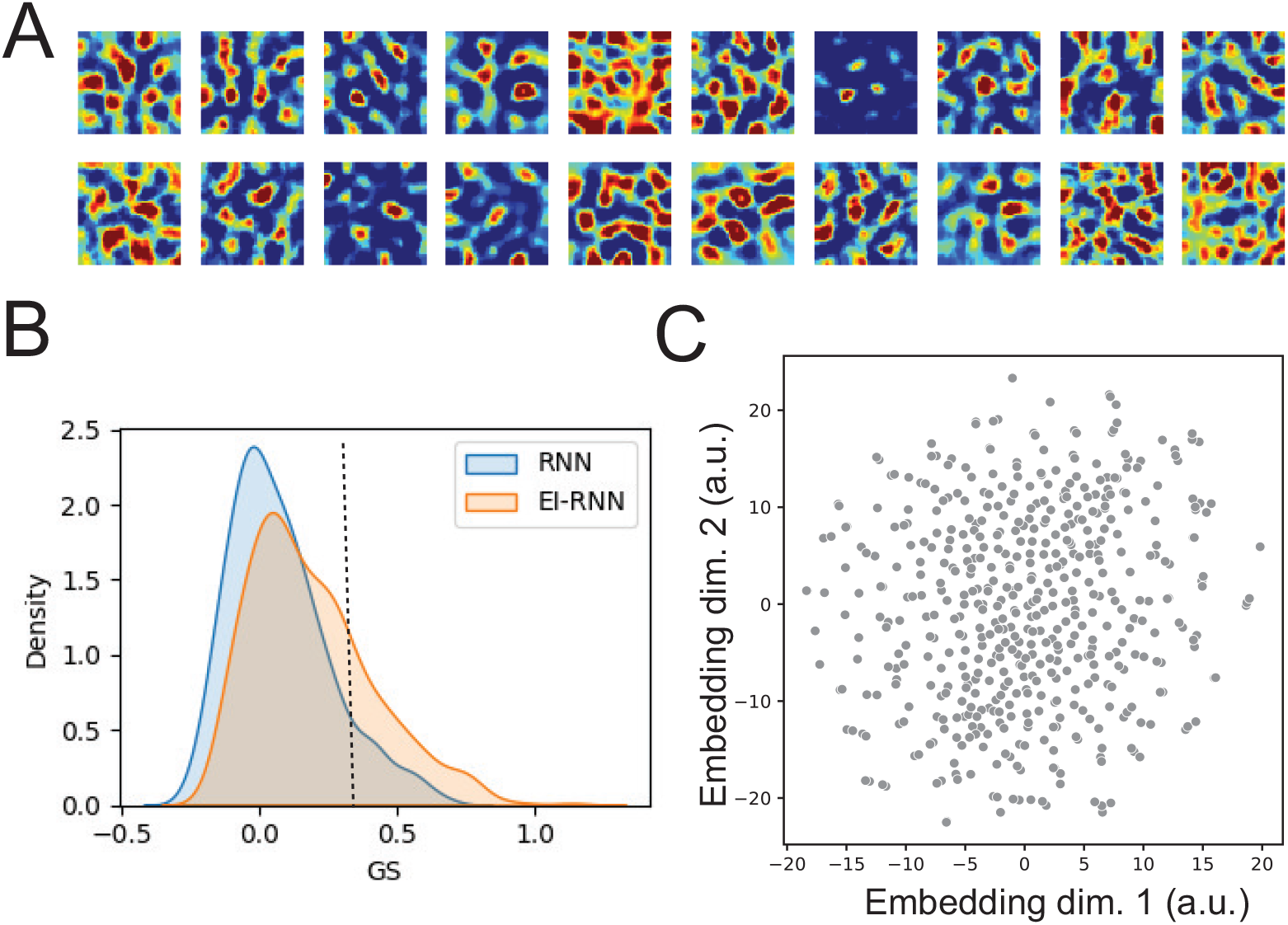
Representative emergent linear RNN unit patterns (Setup #2) without the Dale’s principle constraint; related to Figure 2. **(A)** Examples of spatial firing patterns with respect to the 2D position. **(B)** Compared to the trained linear E/I-RNN, the grid score (GS) distribution comparison showed a reduced number of grid-like unit patterns. The distributions were estimated by a smooth kernel density estimator. Vertical line indicates the 0.3 threshold for grid units. **(C)** 2D embedding of 512 learned unit patterns from the trained linear RNN. No clear clustered structure was found in the embedding space.

**Figure S3:**
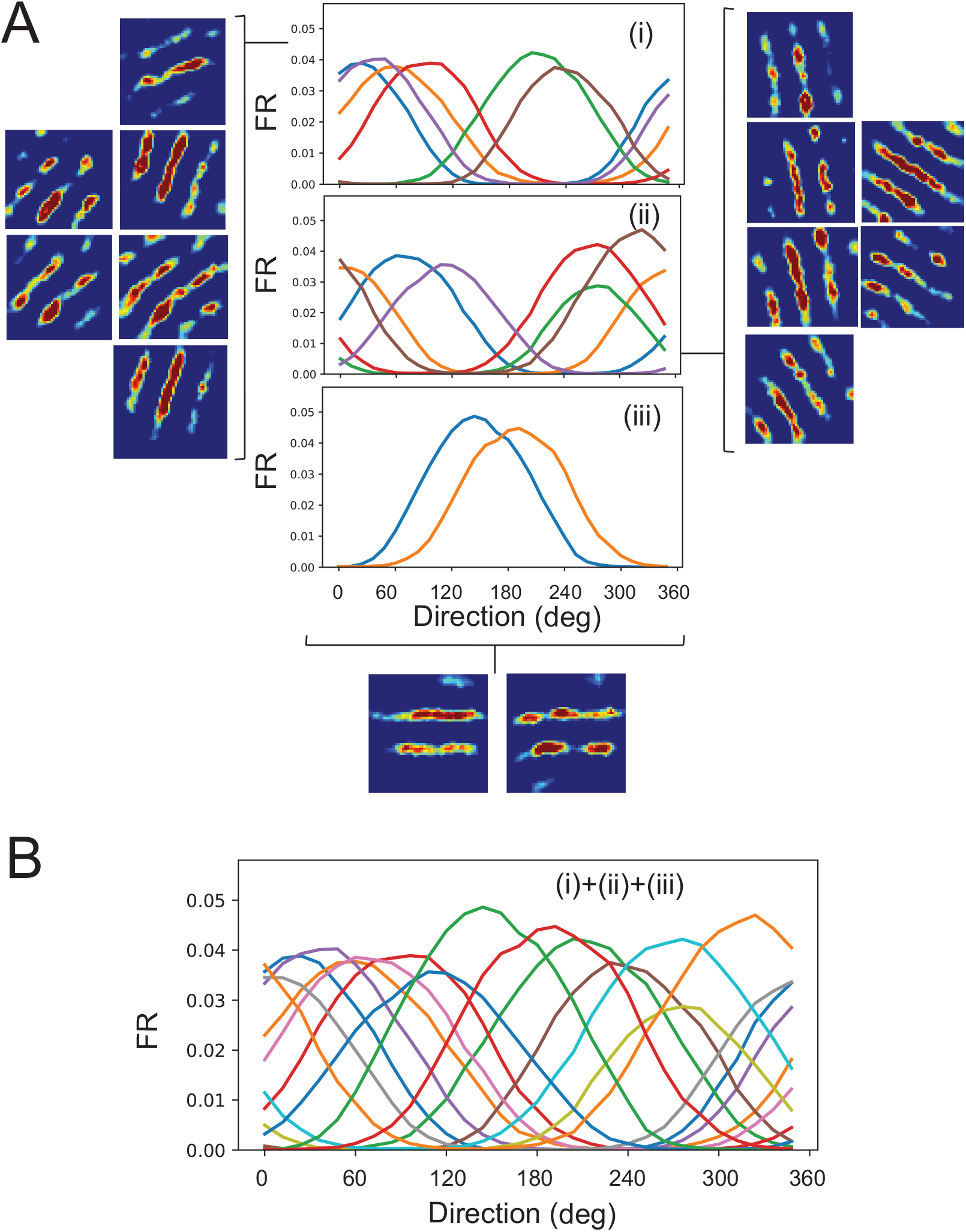
Directional tuning of band-like units; related to Figure 3. **(A)** Many band-like patterns displayed cosine-shaped direction tuning (Setup #2). Classes (i), (ii) and (iii) had different preferences in direction tuning. **(B)** Population of band-like units formed an approximately uniform coverage in direction tuning between 0° and 360°.

**Figure S4:**
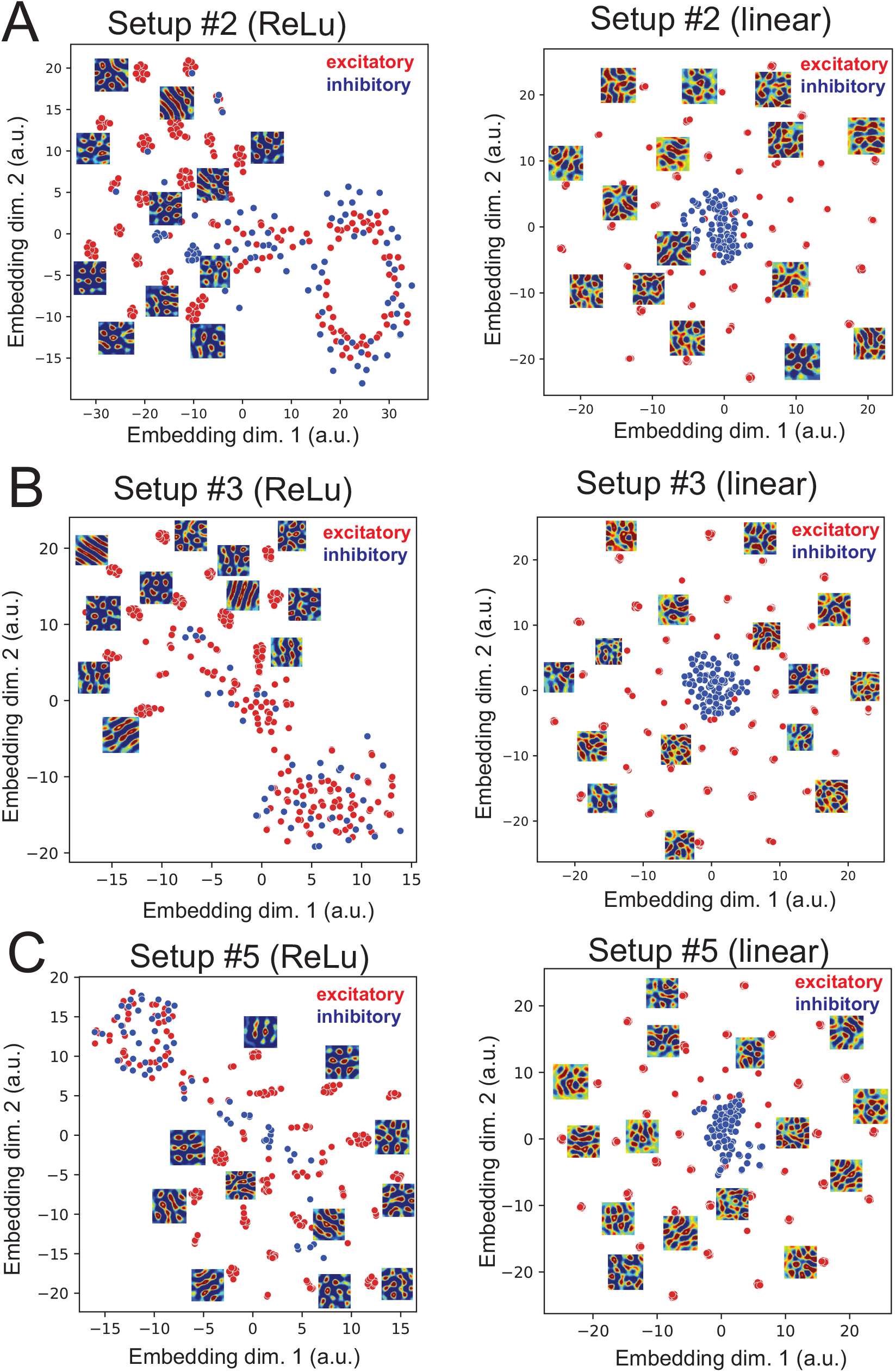
2D embedding of unit firing patterns from the trained E/I-RNNs; related to Figure 3. **(A)** Setup #2 (left: ReLu; right: linear unit), similar to Figure 3E. **(B)** Setup #3 (left: ReLu; right: linear unit). **(C)** Setup #5 (left: ReLu; right: linear unit).

**Figure S5:**
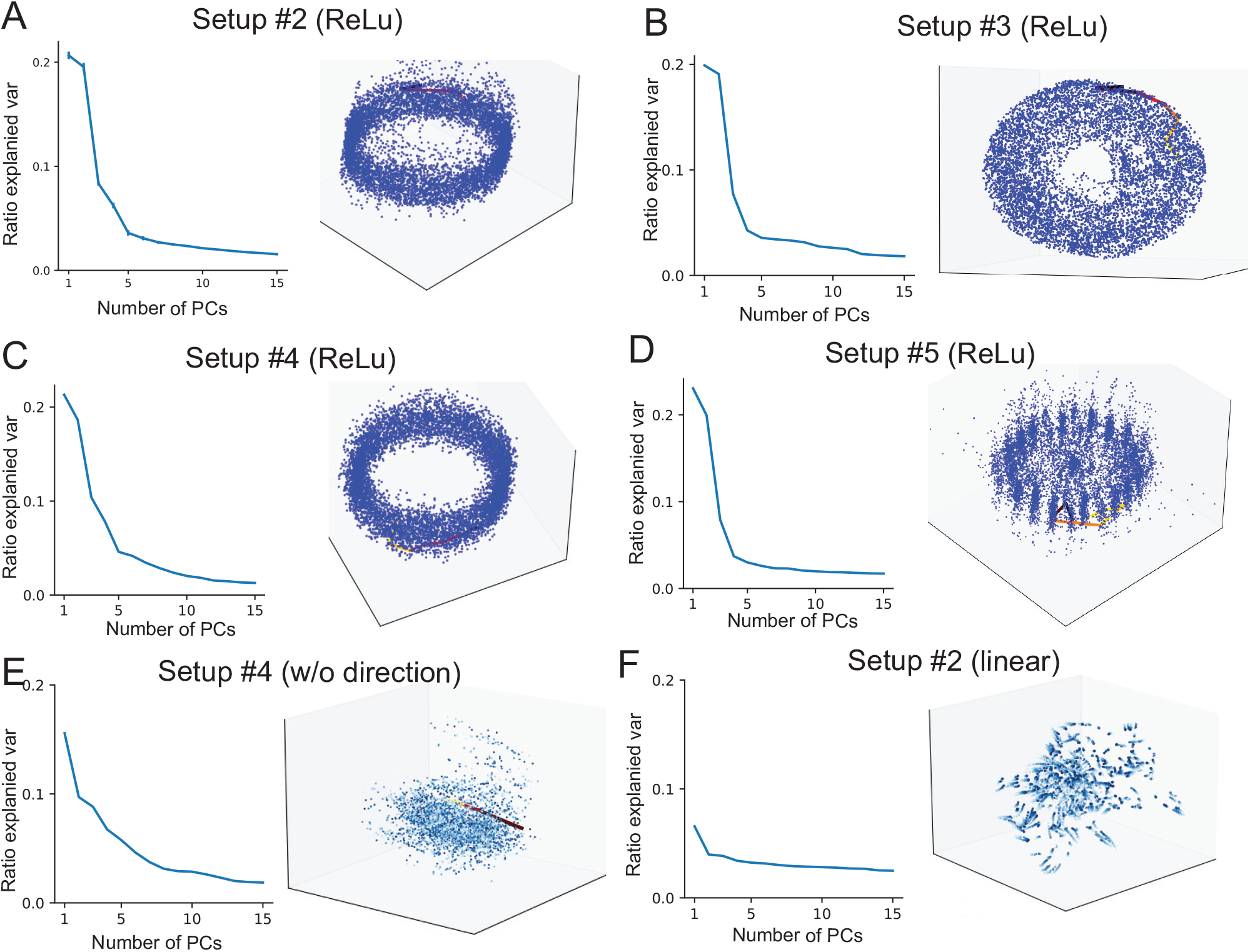
Explained variance of PCA and associated 3D manifolds for various E/I- RNN input configurations and activations; related to Figure 4. **(A)** E/I-RNN with ReLu activation function (Setup #2). **(B)** E/I-RNN with ReLu activation function (Setup #3). **(C)** E/I-RNN with ReLu activation function (Setup #4). **(D)** E/I-RNN with ReLu activation function (Setup #5). **(E)** E/I-RNN with ReLu activation function (Setup #4 but without the direction input). **(F)** E/I-RNN with linear activation function (Setup #2).

**Figure S6:**
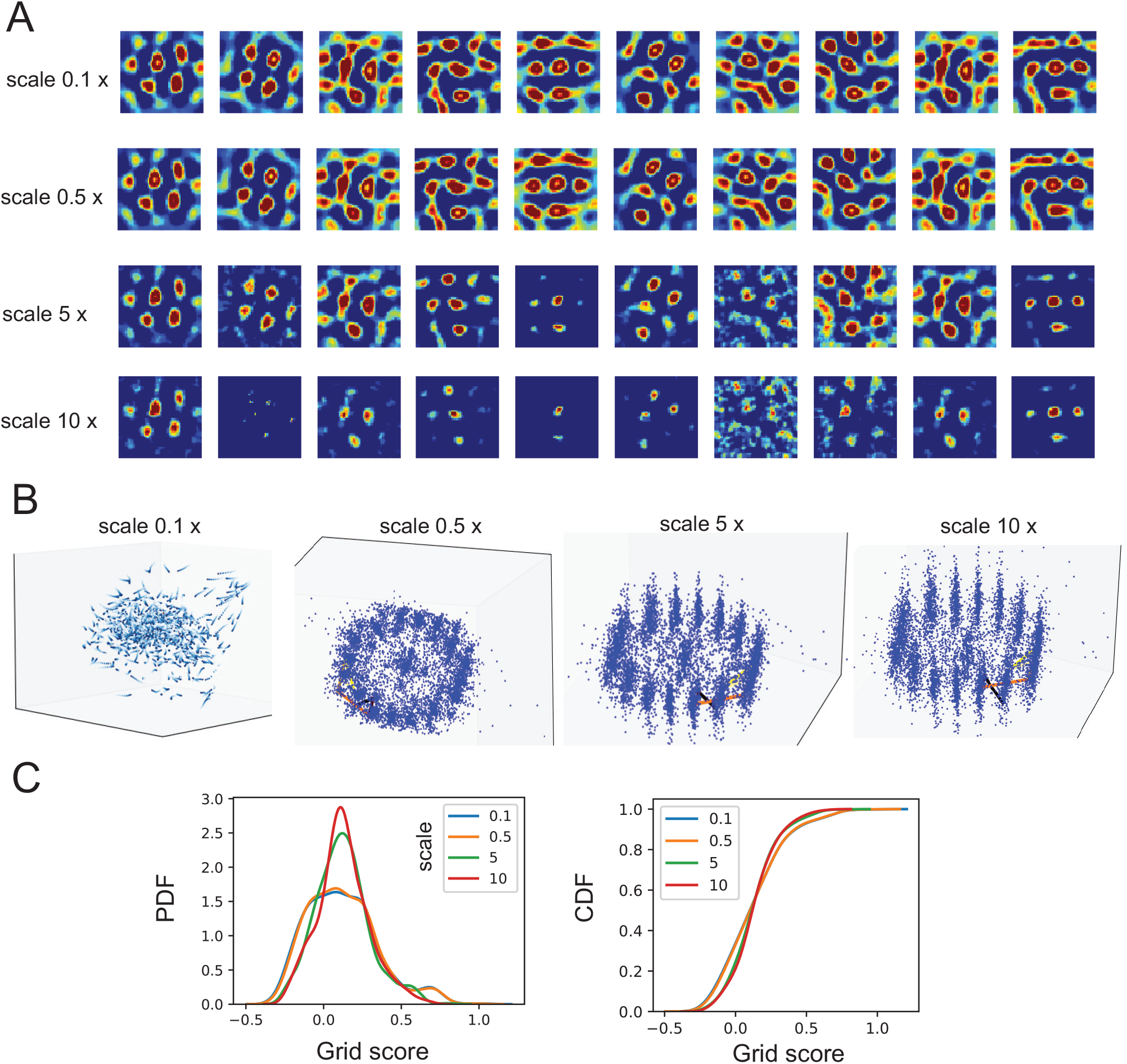
Robustness of grid patterns with respect to the speed of visual optical flow (Setup #5); related to Figure 5. **(A)** Comparison of grid unit patterns when the speed scale of optical flow was varied (baseline: scale 1). **(B)** The corresponding 3D manifolds of population responses under different scales. **(C)** Comparison of GS statistics in probability density distribution (PDF) and cumulative distribution function (CDF) under different scales.

**Figure S7:**
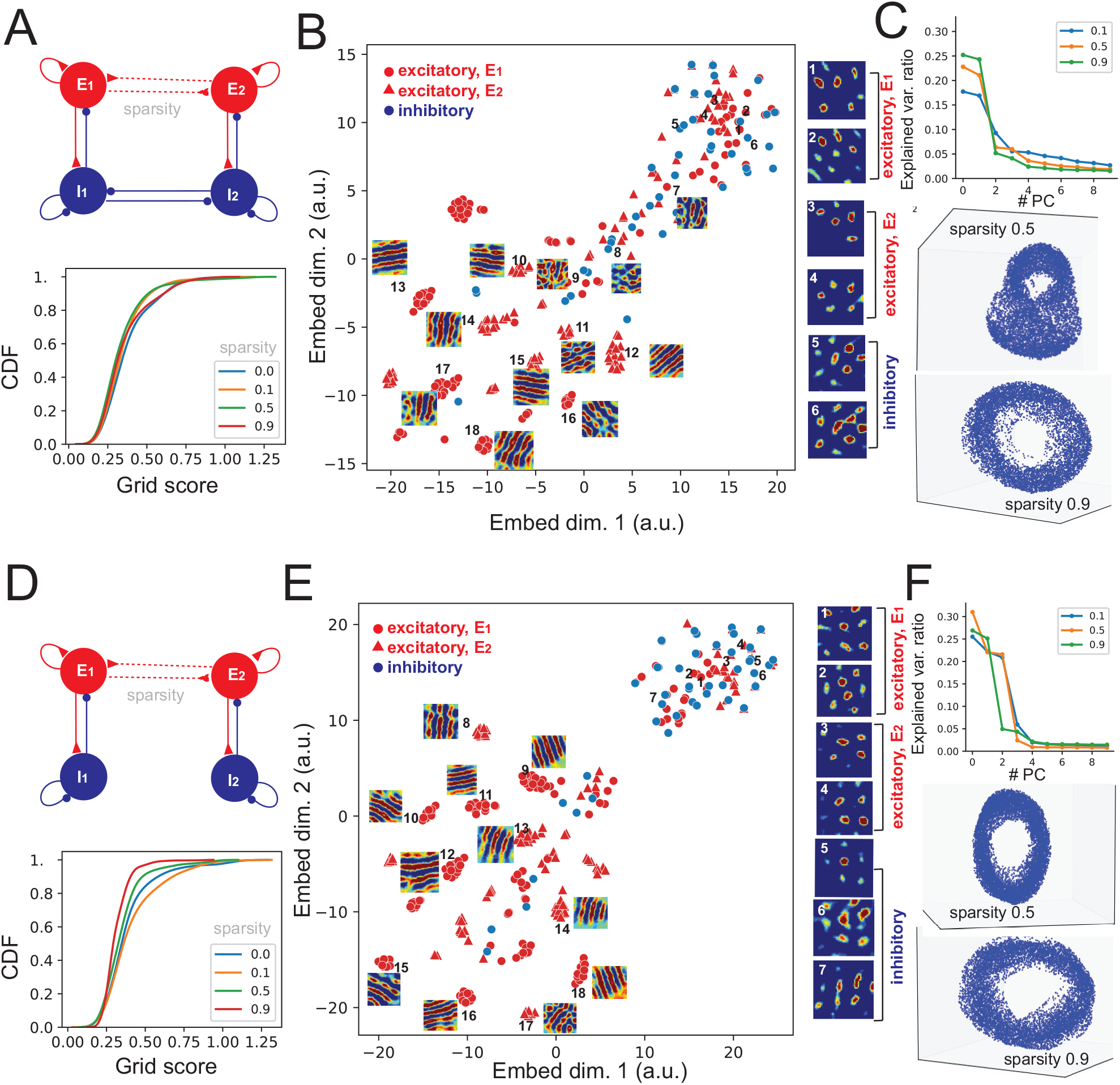
Grid patterns produced by E/I-RNNs with different subnetwork connec- tivity; related to Figure 6. **(A)** Grid score distribution statistics for the selected subnetwork connectivity (top panel). The dashed line denotes weak connections with various sparsity levels (0, 0.1, 0.5, 0.9). Sparsity level 0 implies full connections (original setting). **(B)** 2D projection of learned grid fields (top 50% grid scores) via the t-SNE algorithm. Representative grid fields are shown from each functional groups. **(C)** PCA explained variance ratio (top) and 3D ring manifold of RNN population responses for two sparsity levels. **(D-F)** Similar to panels A-C, except for a different subnetwork connectivity.

**Figure S8:**
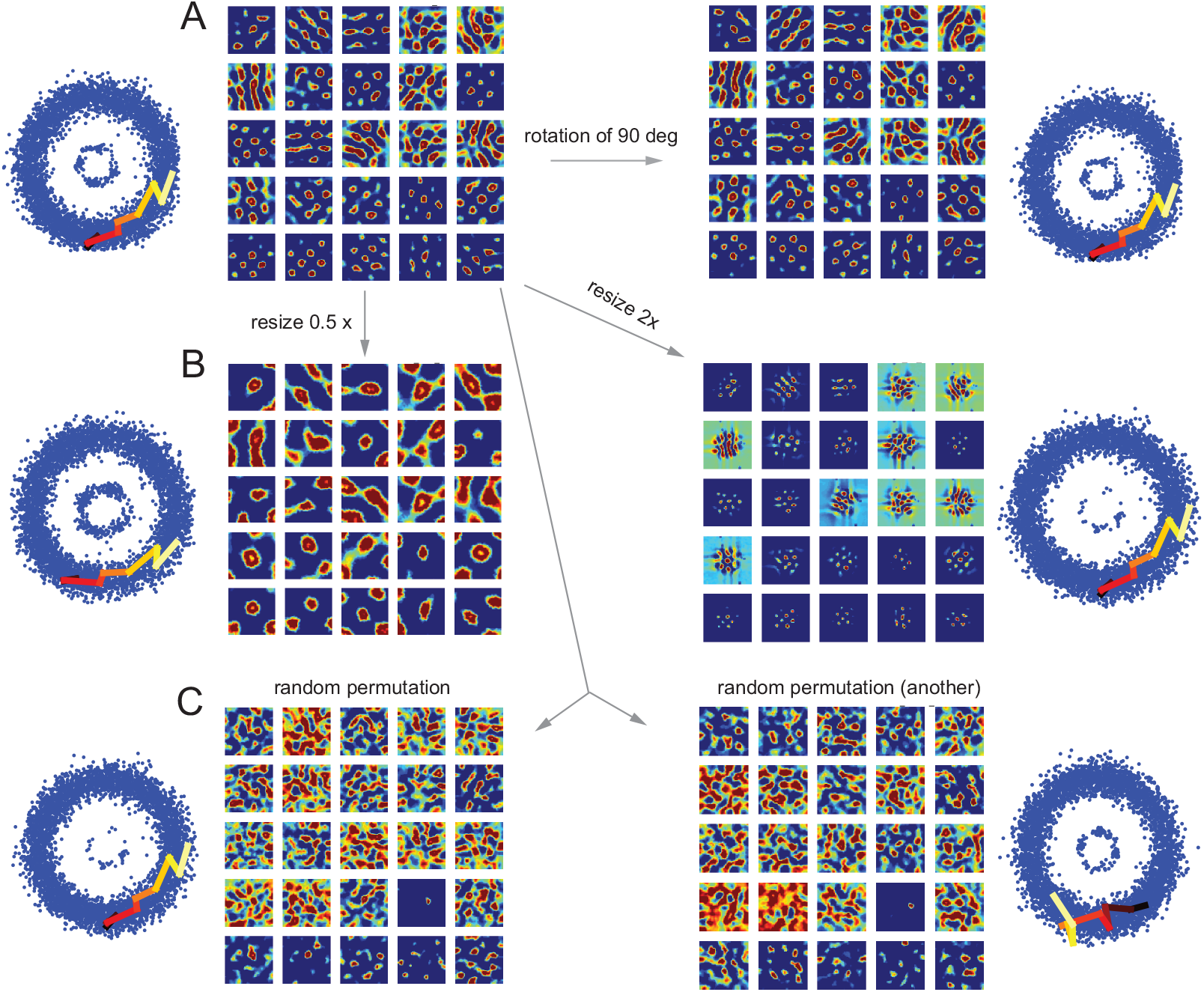
Remapping of visual grid patterns induced by the changes in output place fields; related to Figure 6. **(A)** Changes in grid patterns as a result of rotation of place fields in the RNN output layer. **(B)** Changes in grid patterns as a result of resizing place fields in the RNN output layer. **(C)** Changes in grid patterns as a result of permutation of place fields in the RNN output layer.

**Figure S9:**
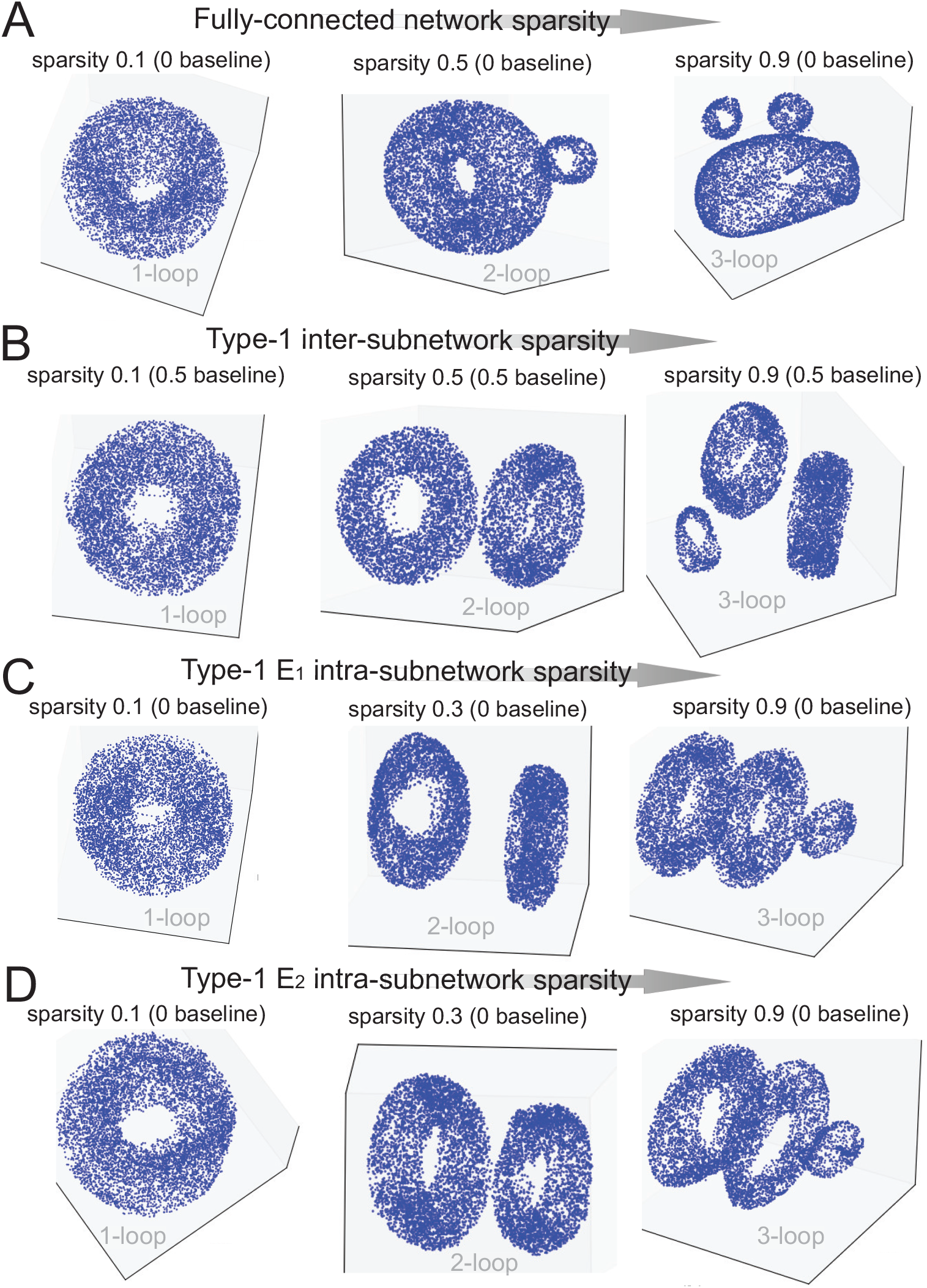
Emerged multistable ring attractors with increasing sparsity in network connectivity; related to Figure 7. **(A)** Evolution of the 3D manifold structure with increasing fully-connected network connectivity (Setup #4, sparsity baseline 0). **(B)** Evolution of the 3D manifold structure with increasing inter-subnetwork connectivity (Type 1, Setup #4, sparsity baseline 0.5). **(C)** Evolution of the 3D manifold structure with increasing *E*_1_ intra-subnetwork connectivity, whereas the *E*_2_ intra-subnetwork connectivity remained unchanged (Type 1, Setup #4, sparsity baseline 0). **(D)** Evolution of the 3D manifold structure with increasing *E*_2_ intra-subnetwork connectivity, whereas the *E*_1_ intra-subnetwork connectivity remained unchanged (Type 1, Setup #4, sparsity baseline 0).

**Figure S10:**
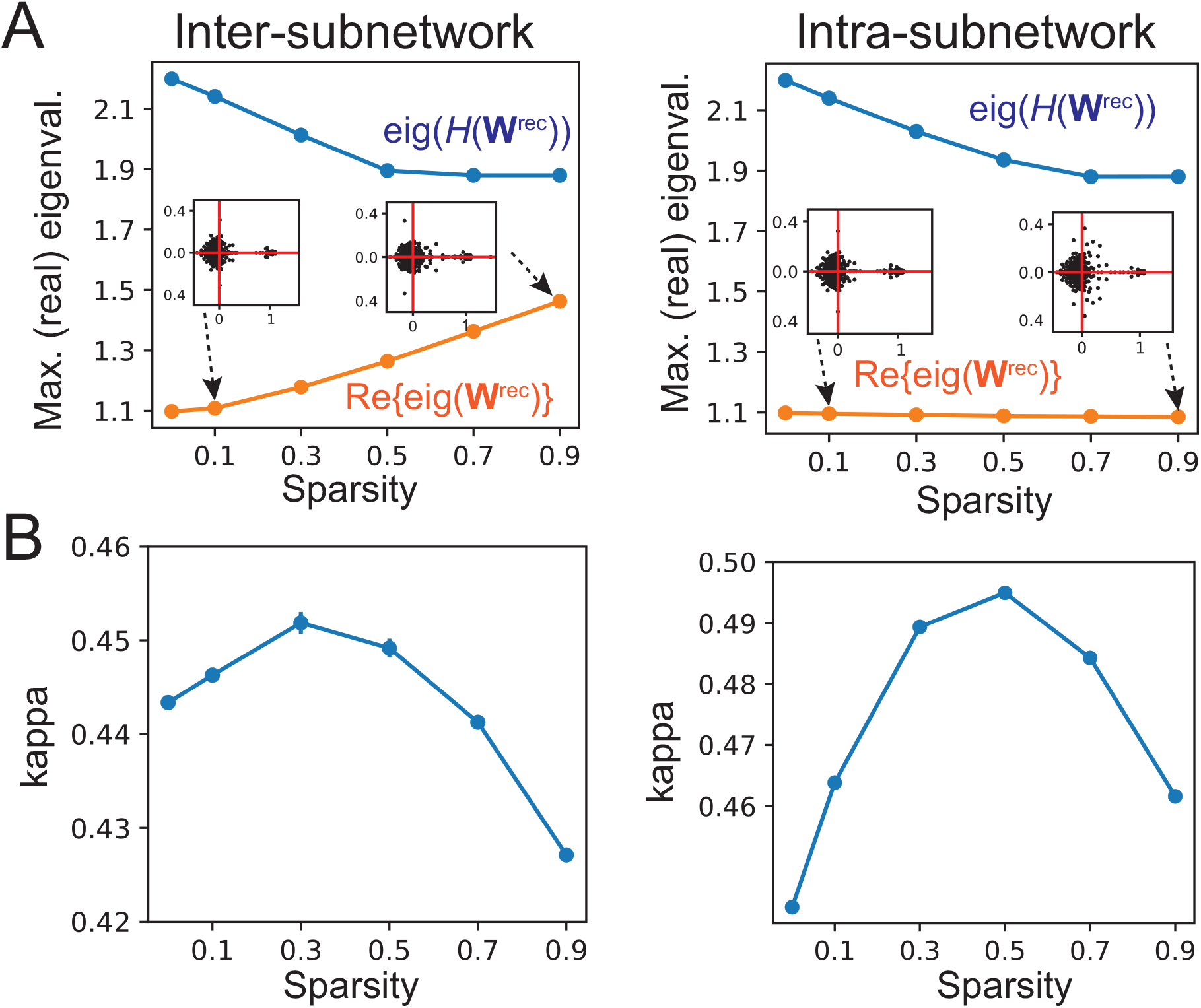
Changes in properties of W^rec^ with respect to the connectivity sparsity of E/I-RNN (Setup #2, Type-1); related to Figure 7. **(A)** Maximum (real part) eigenvalues of of **W**^rec^ and 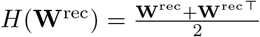 are shown with respect to the sparsity level. The two inset panels represent the eigenvalue spectra at sparsity level 0.1 and 0.9. **(B)** Strength (*κ*) of functionally feedforward connections (STAR Methods) with respect to the sparsity level. All error bars denote SD from 10 random realizations. Sparsity 0 represents the baseline condition.

## References

Alexander AS, Carstensen LC, Hinman JR, et al. (2020). Egocentric boundary vector tuning of the retrosplenial cortex. Science Advances 6, eaaz2322.

Agmon H, Burak Y (2020). A theory of joint attractor dynamics in the hippocampus and the entorhinal cortex accounts for artificial remapping and grid cell field-to-field variability. eLife 9, e56894.

Asllani M, Lambiotte R, Carletti T (2018). Structure and dynamical behavior of non-normal networks. Science Adv. 4(12), sciadv.aau9403.

Averna A, Pasquale V, Murphy MD, et al. (2020). Differential effects of open- and closed-Loop intracortical microstimulation on firing patterns of neurons in distant cortical areas. Cereb Cortex 30, 2879–2896.

Averna A, Hayley P, Murphy MD, et al. (2021). Entrainment of network activity by closed-loop microstimulation in healthy ambulatory rats. Cereb Cortex 31, 5042–5055.

Banino A, Baary C, Uria B, et al. (2018). Vector-based navigation using grid-like representations in artificial agents. Nature, 557, 429–433.

Bao X, Gjorgieva E, Shanahan LK, et al. (2019). Grid-like neural representations support olfactory navigation of a two-dimensional odor space. Neuron, 102, 1066–1075.

Barry C, Ginzberg LL, O’Keefe, J. and Burgess, N. (2012). Grid cell firing patterns signal environmental novelty by expansion. Proc. Natl. Acad. Sci. USA 109, 17687–17692.

Becht E, McInnes L, Healy J, Dutertre C-A, et al. (2019). Dimensionality reduction for visualizing single-cell data using UMAP. Nat. Biotech. 37, 38–44.

Behrens TEJ, Muller TH, Whittington JCR, et al. (2018). What is a cognitive map? Organizing knowledge for flexible behavior. Neuron, 100, 490–509.

Bellmund JL, Deuker L, Schroder TN, Doeller CF (2018a). Grid-cell representations in mental stimulation. eLife, 5, e17089.

Bellmund JL, Gardenfors P, Moser EI, Doeller CF (2018b). Navigating cognition: spatial codes for human thinking. Science 362, eaat6766.

Bicanski A and Burgess N (2019). A Computational model of visual recognition memory via grid cells. Curr Biol 29, 979–990.

Brandon, M. P., Bogaard, A. R., Libby, C. P., Connerney, M. A., Gupta, K. and Hasselmo, M. E. (2011). Reduction of theta rhythm dissociates grid cell spatial periodicity from directional tuning. Science 332, 595–599.

Bridi MCD, Zong FJ, Min X, et al. (2020). Daily oscillation of the excitation-inhibition balance in visual cortical circuits. Neuron, 105, 621–629.

Burak Y, Fiete IR (2009). Accurate path integration in continuous attractor network models of grid cells. PLoS Comput. Biol. 5, e1000291.

Burak Y (2014). Spatial coding and attractor dynamics of grid cells in the entorhinal cortex. Curr. Opin. Neurobiol. 25, 169–175.

Burgess N, Barry C, O’Keefe J (2007). An oscillatory interference model of grid cell firing. Hippocampus, 17, 801–812.

Burgess N (2008). Grid cells and theta as oscillatory interference: theory and predictions. Hippocampus, 18, 1157–1174.

Bush, D. and Burgess, N. (2014). A hybrid oscillatory interference/continuous attractor network model of grid cell firing. J. Neurosci. 34, 5065–5079.

Bush, D., Barry, C., and Burgess, N. (2014). What do grid cells contribute to place cell firing? Trends Neurosci. 37, 136–145.

Bush, D., Barry, C., Manson, D. and Burgess, N. (2015). Using grid cells for navigation. Neuron 87, 507–520.

Buxhoeveden, D. P. (2002). The minicolumn hypothesis in neuroscience. Brain. 125, 935–951.

Campbell MG, Giocomo LM (2018). Self-motion processing in visual and entorhinal cortices: inputs, integration, and implications for position coding. J. Neurophysiol. 120, 2091–2106.

Cao L, Varga V, Chen ZS (2021) Uncovering spatial representations from spatiotemporal patterns of rodent hippocampal field potentials. Cell Rep. Methods, 1, 100101.

Chen, G., King, J.A., Burgess, N., and O’Keefe, J. (2013). How vision and movement combine in the hippocampal place code. Proc. Natl. Acad. Sci. USA 110, 378–383.

Chen, G., Manson D, Cacucci F, Wills TJ (2016). Absence of visual input in the disruption of grid cell firing in the mouse. Curr Biol 26, 2335–2342.

Chen, G., Lu, Y., King, J.A. et al. (2019). Differential influences of environment and self-motion on place and grid cell firing. Nat Commun 10, 630.

Cheung, A., Ball, D., Milford, M., Wyeth, G. and Wiles, J. (2012). Maintaining a cognitive map in darkness: the need to fuse boundary knowledge with path integration. PLoS Comput. Biol. 8, e1002651.

Constantinescu AO, O’Reilly JX, Behrens TEJ (2016). Organizing conceptual knowledge in humans with a gridlike code. Science 352, 1464–1468.

Cousey JJ, Wittoelar A, Zhang S-J, et al. (2013). Recurrent inhibitory circuitry as a mechanism for grid formation. Nat. Neurosci. 16, 318–324.

Cueva CJ, Wei XX (2018). Emergence of grid-like representations by training recurrent neural networks to perform spatial localization. Proc. ICLR (https://arxiv.org/abs/1803.07770)

Dannenberg H, Lazaro H, Nambiar P, Hoyland A, Hasselmo ME (2020). Effects of visual inputs on neural dynamics for coding of location and running speed in medial entorhinal cortex. eLife, 9, e62500.

Dayan P (1993) Improving generalization for temporal difference learning: the successor representation. Neural Comput 5, 613–624.

De Pasquale, R., and Sherman, S. M. (2013). A modulatory effect of the feedback from higher visual areas to V1 in the mouse. J. Neurophysiol. 109, 2618–2631.

Diamanti EM, Reddy CB, Schroder S, et al. (2021). Spatial modulation of visual responses arises in cortex with active navigation. eLife, 10, e63705.

Doeller, C.F., Barry, C., and Burgess, N. (2010). Evidence for grid cells in a human memory network. Nature 463, 657–661.

Fiser, A., Mahringer, D., Oyibo, H.K., Petersen, A.V., Leinweber, M., and Keller, G.B. (2016). Experience-dependent spatial expectations in mouse visual cortex. Nature Neuroscience 19, 1658–1664.

Flossmann T, Rochefort NL (2021). Spatial navigation signals in rodent visual cortex. Curr. Opin. Neurobiol. 67, 163–173.

Fournier, J., Saleem, A.B., Diamanti, E.M., Wells, M.J., Harris, K.D., and Carandini, M. (2020). Mouse visual cortex is modulated by distance traveled and by theta oscillations. Current Biology 30, 3811–3817.

Fuhs, M. C. and Touretzky, D. S. (2006) A spin glass model of path integration in rat medial entorhinal cortex. J. Neurosci. 26, 4266–4276.

Fyhn M, Hafting T, Treves A, Moser MB, Moser EI (2007). Hippocampal remapping and grid realignment in entorhinal cortex. Nature 446,190–194.

Gardner RJ, Hermansen E, Pachitariu M, Burak Y, et al. (2022). Toroidal topology of population activity in grid cells. Nature, https://doi.org/10.1038/s41586-021-04268-7

Gershman SJ (2018). The successor representation: Its computational logic and neural substrates. J. Neuroscience 38, 7193–7200;

Ginosar, G., Aljadeff, J., Burak, Y. et al. (2021) Locally ordered representation of 3D space in the entorhinal cortex. Nature 596, 404–409.

Giocomo LM, Moser M-B, Moser EI (2011). Computational models of grid cells. Neuron, 71, 589–603.

Goldman, M. S. (2009). Memory without feedback in a neural network. Neuron 61, 621–634.

Grieves, R.M., Jedidi-Ayoub, S., Mishchanchuk, K. et al. (2021). Irregular distribution of grid cell firing fields in rats exploring a 3D volumetric space. Nat Neurosci 24, 1567–1573.

Hafting T, Fyhn M, Molden S, Moser MB, Moser EI (2005). Microstructure of a spatial map in the entorhinal cortex. Nature 436, 801–806.

Haggerty DC, Ji D (2015). Activities of visual cortical and hippocampal neurons co-fluctuate in freely moving rats during spatial behaviors. eLife, 4, e08902.

Hardcastle K, Maheswaranathan N, Ganguli S, Giocomo LM (2017). A multiplexed, heterogeneous, and adaptive code for navigation in medial entorhinal cortex. Neuron 94, 375–387.

Hennequin G, Vogels TP, Gerstner W (2012). Non-normal amplification in random balanced neuronal networks Phys. Rev. E 86, 011909.

Hok, V., Jacob, P.-Y., Bordiga, P., Truchet, B., Poucet, B., and Save, E. (2018). A spatial code in the dorsal lateral geniculate nucleus. bioRxiv, 473520.

Horn, B. K. P. and Schunck, B. G. (1981) Determining optical flow. Artif. Intell. 17, 185–203.

Horton JC, Adams DL (2005). The cortical column: a structure without a function.Philo. Trans. R. Soc. Lond. Biol. Sci. 360, 837–862.

Jacobs J, Weidemann CT, Miller JF, et al. (2013). Direct recordings of grid-like neuronal activity in human spatial navigation. Nat. Neurosci. 16, 1188–1190.

Ji D, Wilson MA (2007). Coordinated memory replay in the visual cortex and hippocampus during sleep. Nat. Neurosci. 10, 100–107.

Kang L, Balasubramanian V (2019). A geometric attractor mechanism for self-organization of entorhinal grid modules. eLife, 8, e46687.

Kerg G, Goyette K, Touzel P, et al. (2019). Non-normal recurrent neural netwok (nnRNN): learning long time dependencies while improving expressive with transient dynamics. Proc. 33rd Conf. Neural Info. Proc. Syst. (NeurIPS’19).

Kim R, Sejnowski TJ (2021). Strong inhibitory signaling underlies stable temporal dynamics and working memory in spiking neural networks. Nat. Neurosci. 24, 129–139.

Klukas M, Lewis M, Fiete I (2020) Efficient and flexible representation of higher-dimensional cognitive variables with grid cells. PLoS Comput Biol 16(4): e1007796.

Krupic J, Brugess N, O’Keefe J (2012). Neural representations of location composed of spatially periodic bands. Science, 337, 853–857.

Laramee ME, Rockland KS, Prince S, Bronchti G, Boire D (2013). Principal component and cluster analysis of Layer V pyramidal cells in visual and non-visual cortical areas projecting to the primary visual cortex of the mouse. Cerebral Cortex, 23, 714–728.

Lappe M, Rauschecker J (1992). Computation of heading direction from optical flow in visual cortex. Advances in Neural Information Processing Systems 5 (NIPS).

Leinweber, M., Ward, D.R., Sobczak, J.M., Attinger, A., and Keller, G.B. (2017). A sensorimotor circuit in mouse cortex for visual flow predictions. Neuron 95, 1420–1432 e1425.

Litwin-Kumar, A., Doiron, B (2012). Slow dynamics and high variability in balanced cortical networks with clustered connections. Nat Neurosci 15, 1498–1505.

Liu L, She L, Chen M, et al. (2016). Spatial structure of neuronal receptive field in awake monkey secondary visual cortex (V2). Proc. Natl. Acad. Sci. USA, 113, 1913–1918.

Long X, Zhang S-J (2021). A novel somatosensory spatial navigation system outside the hippocampal formation. Cell Res., 31, 649–663.

Long X, Deng B, Cai J, Chen ZS, Zhang S-J (2021). A compact spatial map in V2 visual cortex. BioRxiv preprint. https://www.biorxiv.org/content/10.1101/2021.02.11.430687v1

Long X, Cai J, Deng B, Chen ZS, Zhang S-J (2021). Bimodal remapping in visual grids. BioRxiv preprint. https://www.biorxiv.org/content/10.1101/2021.10.30.466568v1.

Long X, Deng B, Young CK, et al. (2022). Sharp tuning of head direction and angular velocity cells in the somatosensory cortex. Advanced Sciences, 2022, 202200020.

Marshel JH, Garrett ME, Nauhaus I and Callaway EM (2011). Functional specialization of seven mouse visual cortical areas. Neuron 72, 1040–1054.

McInnes, L, Healy, J (2018). UMAP: Uniform Manifold Approximation and Projection for Dimension Reduction. ArXiv e-prints 1802.03426.

McNaughton, B. L., Battaglia, F. P., Jensen, O., Moser, E. I., Moser, M.-B. (2006). Path integration and the neural basis of the “cognitive map”. Nat. Rev. Neurosci. 7, 663–678.

Michaels JA, Dann B, Scherberger H (2016). Neural population dynamics during reaching are better explained by a dynamical system than representational tuning. PLoS Comput. Biol. 12, e1005175.

Miller MW, Vogt BA (1984). Direct connections of rat visual cortex with sensory, motor, and association cortices. J Comp Neurol. 226, 184–202.

Mok, R.M., Love, B.C. (2019). A non-spatial account of place and grid cells based on clustering models of concept learning. Nat Commun 10, 5685.

Murphy BK and Miller KD (2009). Balanced amplification: a new mechanism of selective amplification of neural activity patterns. Neuron 61, 635–648.

Najafi F, Elsaye GF, Gao R, Pnevmatikakis E, et al. (2020). Excitatory and inhibitory subnetworks are equally selective during decision-making and Emerge simultaneously during learning. Neuron, 105, 165–179.

Nau M, Schroder TNB, Bellmund JLS, Doeller CF (2018). Hexadirectional coding of visual space in human entorhinal cortex. Nat Neurosci 21, 188–190.

Narvatilova Z, Godfrey KB, McNaughton BL (2016). Grids from bands, or bands from grids? An examination of the effects of single unit contamination on grid firing patterns. J. Neurophysiol. 115, 992–1002.

Orchard J, Yang H, Ji X (2013). Does the entorhinal cortex use the Fourier transform? Front. Comput. Neurosci. 7, 179.

Patra M (2018). Multiple attractor bifurcation in three-dimensional piecewise linear maps. International Journal of Bifurcation and Chaos, 28, 1830032.

Pollock E, Jazayeri M (2020) Engineering recurrent neural networks from task-relevant manifolds and dynamics. PLoS Comput Biol 16, e1008128.

Raithel CU, Gottfried JA (2021). What are grid-like responses doing in the orbitofrontal cortex? Behav. Neurosci. 135, 218–225.

Rajakumar A, Rinzel J, Chen ZS (2021). Stimulus-driven and spontaneous dynamics in excitatory-inhibitory recurrent neural networks for sequence representation. Neural Computation, 33, 2603–2645.

Recanatesi S, Farrell M, Lajoie G, Deneve S, Rigotti M, Shea-Brown E (2021). Predictive learning as a network mechanism for extracting low-dimensional latent space representations. Nat. Comm. 12, 1417.

Rosay S, Weber S, Mulas M (2019). Modeling grid fields instead of modeling grid cells. J. Comp. Neurosci. 47, 43–60.

Rowland DC, Roudi Y, Moser MB, Moser EI (2016). Ten years of grid cells. Annu. Rev. Neurosci. 39, 19–40.

Rueckemann, J.W., Sosa, M., Giocomo, L.M., Buffalo, E.A. (2021). The grid code for ordered experience. Nat. Rev. Neurosci. 22, 637–649.

Saleem AB, Diamanti EM, Fournier J, et al. (2018). Coherent encoding of subjective spatial position in visual cortex and hippocampus. Nature, 562, 124–127.

Sanderson, K.J., Dreher, B., Gayer, N. (1991). Prosencephalic connections of striate and extrastriate areas of rat visual cortex. Exp Brain Res 85, 324–334.

Savelli F and Knierim JJ (2019). Origin and role of path integration in the cognitive representations of the hippocampus: computational insights into open questions. J Exp Biol (2019) 222, jeb188912.

Seung HS (1998). Continuous attractors and oculomotor control. Neural Networks, 11, 1253–1258.

Shilnikov AL, Maurer AP (2016). The art of grid fields: geometry of neuronal time. Front. Neural Circuits, 10, 12.

Song, H.F., Yang, G.R., and Wang, X.-J. (2016). Training excitatory-inhibitory recurrent neural networks for cognitive tasks: A simple and flexible framework. PLoS Comput. Biol. 12, e1004792.

Song Z, Xu J, Zhen B (2019). Mixed-coexistence of periodic orbits and chaotic attractors in an inertial neural system with a nonmonotonic activation function. Mathematical Biosciences and Engineering, 6, 6406–6425.

Sorscher B, Mel GC, Ocko SA, Giocomo L, Ganguli S (2020). A unified theory for the computational and mechanistic origins of gird cells. https://www.biorxiv.org/content/10.1101/2020.12.29.42483v1.

Stachenfeld, K., Botvinick, M, Gershman, S. (2017). The hippocampus as a predictive map. Nat. Neurosci 20, 1643–1653.

Sussillo, D. and Abbott, L.F. (2009). Generating coherent patterns of activity from chaotic neural networks. Neuron, 63, 544–557.

Sussillo, D., Barak, O. (2013) Opening the black box: low-dimensional dynamics in high-dimensional recurrent neural networks. Neural Computation 25, 626–649.

Sussillo, D., Churchland, M.M., Kaufman, M.T., and Shenoy, K. V. (2015). A neural network that finds a naturalistic solution for the production of muscle activity. Nature Neuroscience, 18, 1025–1033.

Taube JS (2007). The head direction signal: origins and sensory-motor integration. Ann. Rev. Neurosci. 30, 181–207.

van der Maaten LJP and Hinton GE. (2008) Visualizing data using t-SNE. Journal of Machine Learning Research, 9, 2579–2605.

Vinepinsky E, Perchik S, Segev R (2020). A generalized linear model of a navigation network. Front. Neural Circuits, 14, 56.

Wang HT, Mathur B, Koch C (1989). Computing optical flow in the primate visual system. Neural Computation, 1, 92–103.

Weber SN, Sprekeler H (2018). Learning place cells, grid cells and invariances with excitatory and inhibitory plasticity. eLife, 7, e34560.

Willmore BDB, Prenger RJ, Gallant JL (2010). Neural representation of natural images in visual area V2. J. Neurosci. 30, 2102–2114.

Wills TJ, Lever C, Cacucci F, Burgess N, O’Keefe J (2005). Attractor dynamics in the hippocampal representation of the local environment. Science, 308, 873–876.

Wurtz RH (1998). Optic flow: a brain region devoted to optic flow analysis? Curr. Biol. 8, 554–556.

Xue, X., Wimmer RD, Halassa, M.M., and Chen, Z.S. (2022). Spiking recurrent neural networks represent task-relevant neural sequences in rule-dependent computation. Cognitive Computation

Yartsev, M. M., Witter, M. P. and Ulanovsky, N. (2011). Grid cells without theta oscillations in the entorhinal cortex of bats. Nature 479, 103–107.

Yao H and Li C-Y (2002). Clustered organization of neurons with similar extra-receptive field properties in the primary visual cortex. Neuron 35, 547–553.

Yu LQ, Park SA, Sweigart SC, Boorman ED, Nassar MR (2021). Do grid codes afford generalization and flexible decision-making? https://arxiv.org/pdf/2106.16219.pdf

Zhang X, Liu S, and Chen ZS (2021). A geometric framework for understanding dynamic information integration in context-dependent computation. iScience 24, 102919.

Zong W, Obenhaus HA, Skytoen ER, et al. (2022). Large-scale two-photon calcium imaging in freely moving mice. Cell, in press. doi.org/10.1016/j.cell.2022.02.017

